# Pervasive Induction of Regulatory Mutation Microclones in Sun-exposed Skin

**DOI:** 10.1101/2024.09.12.612526

**Authors:** Vijay Menon, Alejandro García-Ruiz, Susan Neveu, Brenda Cartmel, Leah M. Ferrucci, Meg Palmatier, Christine Ko, Kenneth Y. Tsai, Mio Nakamura, Sa Rang Kim, Michael Girardi, Karl Kornacker, Douglas E. Brash

## Abstract

Carcinogen-induced mutations are thought near-random, with rare cancer-driver mutations underlying clonal expansion. Using high-fidelity Duplex Sequencing to reach a mutation frequency sensitivity of 4×10^-9^ per nt, we report that sun exposure creates pervasive mutations at sites with ∼100-fold UV-sensitivity in RNA-processing gene promoters – cyclobutane pyrimidine dimer (CPD) hyperhotspots – and these mutations have a mini-driver clonal expansion phenotype. Numerically, human skin harbored 10-fold more genuine mutations than previously reported, with neonatal skin containing 90,000 per cell; UV signature mutations increased 8,000-fold in sun-exposed skin, averaging 3×10^-5^ per nt. Clonal expansion by neutral drift or passenger formation was nil. Tumor suppressor gene hotspots reached variant allele frequency 0.1-10% via 30-3,000 fold clonal expansion, in occasional biopsies. CPD hyperhotspots reached those frequencies in every biopsy, with modest clonal expansion. In vitro, tumor hotspot mutations arose occasionally over weeks of chronic low-dose exposure, whereas CPD hyperhotspot mutations arose in days at 1000-fold higher frequencies, growing exponentially. UV targeted mini-drivers in every skin cell.

## Introduction

Genome sequencing revealed that point mutations are far more frequent in normal tissue than expected, prompting increasing interest in their role in the earliest stages of cancer and aging (*1–6*). Early events – key to substituting prevention for treatment – are generally thought of as stochastic, with occasional mutations coming to prominence when their phenotype is selected on. Skin is the best understood tissue with regard to these events: the carcinogen is known to be sunlight, with different dose responses for squamous cell carcinoma, basal cell carcinoma, and melanoma (*7–12*); the primary mutagenic DNA photoproduct is the cyclobutane pyrimidine dimer (CPD) joining two adjacent pyrimidines (*13, 14*); the principal mutation in tumors and precancers is the UV signature mutation of C→T at dipyrimidine sites, most distinctively CC→TT (*1, 15–18*); key tumor suppressor and oncogenes such as *TP53*, *PTCH*, *NOTCH*, and *RAC1* contain these mutations (*15, 17, 19, 20*); phenotypic effects of these mutations are known, such as allowing further UV exposure to drive apoptosis, altered cell differentiation, and precancerous clonal expansion (*1, 21–25*); and protective mechanisms such as melanin shielding (*26, 27*) and nucleotide excision repair are well understood (*28*). Sunlight is also associated with oxidative damage, generated by UV-induced enzymes, photooxidation of melanin, and the endogenous melanin synthesis pathway (*26, 29–31*).

DNA lesions and mutations are generally considered to be rare stochastic events, with hotspots mutated only ∼5 fold more frequently than surrounding sites (e.g., ref. (*13*). In sun-exposed skin, the published mutation frequency (MF_maxI_, including any recurrences of the same mutation; equivalently, “mutant fraction” (*32*)) in oncogenes and tumor suppressor genes is ∼4 per Mb after clonal expansion (*3, 4, 33–35*); this figure lies at the high-dose end of the 10^-7^–10^-5^ frequency of independent new mutations (MF_minI_) extrapolated from classic drug-selection assays on tissue culture cells given a single UV exposure (*32*). The skin figure will include not only the accumulation over years, facilitated by the lower lethality of sunlight exposures than lab exposures, but also the clonal expansion of mutants and the greater number of nucleotides (nt) assayable by sequencing than by selectable phenotype. Mutations in normal skin might serve as genomic dosimeters, a historical record of an individual’s past UV exposure. Particularly sensitive might be CPD hyperhotspots – dinucleotides with the highest CPD induction frequencies in the genome and lying outside the main Poisson distribution – which have up to 500-fold sensitivity to CPD formation and located in promoters of RNA processing genes (*36, 37*). These sites were originally noticed as ETS-1 transcription factor binding sites mutated in melanomas (*38, 39*) and hyperhotspot-adjacent nt were subsequently found to have elevated CPD frequencies (*40, 41*). At least one of these hyperhotspot sites has a phenotype when mutated, possessing elevated promoter activity in a reporter carrying the UV-signature mutation (*38*).

Investigating the origin of mutations in human skin *in vivo* is a formidable challenge because reporter genes are not available, founder mutants are not distinct colonies on a dish, and immunohistochemistry detects mutant clones only for *TP53* (*2, 21, 25*). Ordinary DNA sequencing methods are limited to a VAF detection limit of ∼0.5%, i.e., very large clones, whereas the VAF expected even at a CPD hyperhotspot is closer to 10^-5^ per nt (*32*). Importantly, even mutations above 0.5% are nearly all spurious, arising from DNA damage on one strand of the initial template (*42*). Indeed, published differences between sun-shielded and sun-exposed skin are only ∼3-fold (*34, 43*). We therefore investigated mutation frequencies in normal human skin at CPD hyperhotspots and in cancer genes, using Duplex Sequencing (*32, 42*). This method separately tags top and bottom strands of the template so that only mutations seen on both strands are counted toward the Duplex Consensus Sequence (DCS). The tradeoff is the need for two rounds of target capture followed by sequencing ∼12 PCR progeny for each DCS. The payoff is the ability to easily detect ∼4×10^-5^ mutant DCSs per nt at a single nucleotide (Variant Allele Frequency, VAF) or, by averaging over a large target, a MF of 10^-8^ per nt or lower.

We find 10-fold higher MF_maxI_ values than reported using standard DNA sequencing; 8000-fold increases in UV signature MF_maxI_ with sunlight exposure, despite only 3-fold changes in total mutations; UV mutagenesis targeted at CPD hyperhotspots and having mutation frequencies exceeding those at cancer genes, little role of neutral drift or passenger formation in clonal expansion, and diverse evidence that CPD hyperhotspots have an *in vivo* clonal expansion phenotype greater than seen in a phenotype-less genome region, making them pervasive UV-targeted mini-drivers.

## Results

### High-fidelity sequencing avoids specious mutation hotspots arising from bioinformatics realignment steps

DNA sequence analysis requires aligning DNA sequence reads to a reference genome; alignment accuracy is critical when detecting rare mutations and identifying mutation hotspots. Because the error-prone nature of standard sequencing methods causes inaccuracies in mapping and variant calling, particularly near insertions and deletions, analysis routinely uses non-stringent genome alignment followed by a realignment step near indel sites before calling variants (*44, 45*). The same protocol is conventionally used in Duplex Sequencing prior to calling variants in the Duplex Consensus Sequence (DCS). However, this should not be necessary because Duplex Sequencing virtually eliminates sequencing errors. The fact that Duplex Sequencing provides independent sequences for top and bottom strands of the parental molecule allowed us to notice that realignment often mapped top and bottom strands to different chromosomes. We therefore developed the present stringent IndelCorrected-DCS aligner (Methods) that selects for unique, full-length alignment of reads within each probed genomic region. **Fig. 1** shows that high fidelity Duplex Sequencing together with precise and accurate IndelCorrected-DCS alignment avoids specious variants and mutation hotspots that are introduced if realignment steps precede variant calling.

**Fig. 1.**
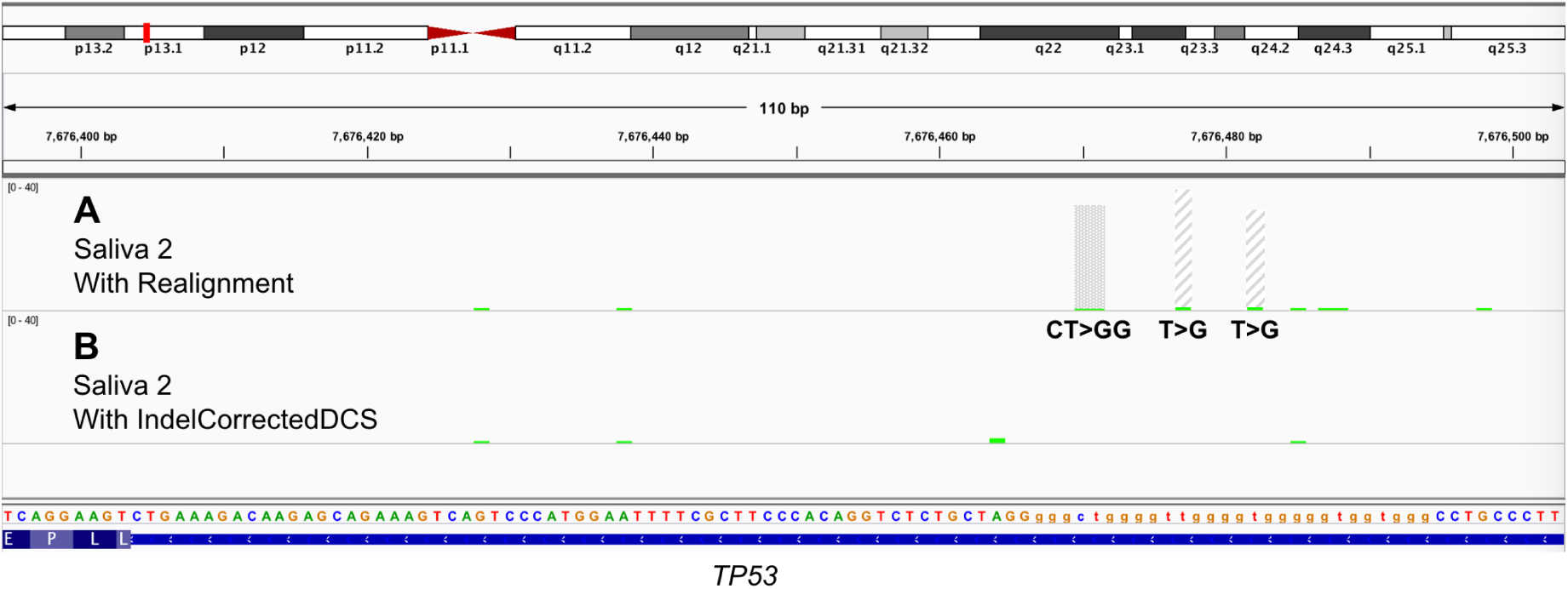
High fidelity sequencing makes it possible to avoid specious variants created when realignment steps precede variant calling. Error-prone sequencing methods and genuine indel mutations cause known inaccuracies in standard mapping procedures and variant calling, prompting routine use of non-stringent genome alignment followed by a realignment step near indel sites before calling variants. In Duplex Sequencing, the same protocol is conventionally used prior to calling variants in the Duplex Consensus Sequence (DCS). Duplex Sequencing virtually eliminates sequencing errors, however, allowing the present stringent IndelCorrected-DCS aligner that selects for unique, full-length alignments of reads within each probed genomic region. This stringency obviates the need for DCS realignment and in fact eliminates errors introduced by realigning DCSs resulting from non-stringent mappings and contributing ∼10% of the apparent sequencing coverage depth. (A) The upper histogram shows standard DCS mutation calls for a region of the *TP53* gene in saliva from an older adult. This chr17 region is G-rich, with a 16 bp indel polymorphism 150 bp 5’ to the region. Inspection of the original alignment files reveals that, at the center site, 39 of the 40 T→G mutations (apparent VAF 2 × 10^-3^) derived from inserts for which the left and right ends had not both mapped to chr17 (crosshatched). The adjacent T→G site had a similar artifact. Reads not in doubt are shown in green. The CT→GG mutation (stippled) was not present in the original alignment files and appeared only after realigned variant calling. (B) The result of stringent initial alignment with omission of realignment. Evidently mis-clustering of rare mutations by conventional procedures creates artifactual mutation hotspots. Even the distinction between low-VAF recurrent sites and count 1 sites is biologically important because the former can reflect clonal expansion and hence phenotypic selection, whereas true count 1 sites reflect independently originating mutations and thus the exposure dose.

### UV-signature VAFs increase from oncogenes to tumor suppressor genes to CPD hyperhotspots

Duplex Sequencing was applied to samples of increasing occasional or frequent sunlight exposure: neonatal foreskin; blood from an 18 year old donor (commercial control); adult buccal epithelia; skin from the leg or forearm having typical ambient sunlight exposure; buttock or forearm of patients therapeutically exposed to nbUVB; and neck skin from a patient with melanoma and squamous cell carcinoma (**Table S1**). Library preparation included two steps of hybrid capture using capture probes for 43-48 specific genome regions that included a commercial low-background control miRNA region (“GenoTox”), presumptively lacking a selectable phenotype (although expressed in brain and testes); oncogenes such as *KRAS* and *BRAF*; tumor suppressor genes such as *TP53* and *NOTCH1*; cancer genes found mutated in sun-exposed skin (*3, 43*); and hyperhotspots for forming UV-induced cyclobutane pyrimidine dimers (CPD) (*36, 37*).

We first focused on the classic UV signature mutation, a C→T substitution at a dipyrimidine (henceforth “C→T diPy”), including the distinctive CC→TT (*18, 46*). These arise at DNA photoproducts involving adjacent T or C nt; in mammalian cells the mutagenic photoproducts are primarily CPDs (*13*) because the other photoproducts are rapidly repaired. **Fig. 2** shows exemplars of C→T mutation distributions in several classes of targets. Background at a nt in neonatal foreskin or blood was 0-1 UV signature mutant DCS per ∼25,000 DCSs (VAF ≤ 4×10^-5^ per nt), whereas non UV signature mutations (mainly T→G and G→T) were present at most nt and at 5-10 fold higher counts. In the GenoTox toxicology monitoring region, UV signature mutation counts reached ∼20-fold higher levels with sun- or UV-exposure. Mutation counts in oncogenes such as *KRAS* and *BRAF* were low, even with sun-exposure, and were not localized to gain-of-function tumor mutation hotspots; they were dropped from the capture panel for later samples. Tumor suppressor genes such as *TP53* and *NOTCH1-3* reached ∼300 counts with UV exposure, primarily at sites mutated in tumors. CPD hyperhotspots, sites at which CPDs arise up to 500-fold more frequently than the genomic average (*36, 37*), were mutated at even higher frequencies, exceeding 1000 counts (i.e., genomes) for some hyperhotspots in some individuals who had only ambient exposure to sunlight. Hyperhotspot mutations were restricted to CPD hyperhotspots observed in keratinocytes although, for some probes, one or two nearby nt were also recurrently mutated.

**Fig. 2.**
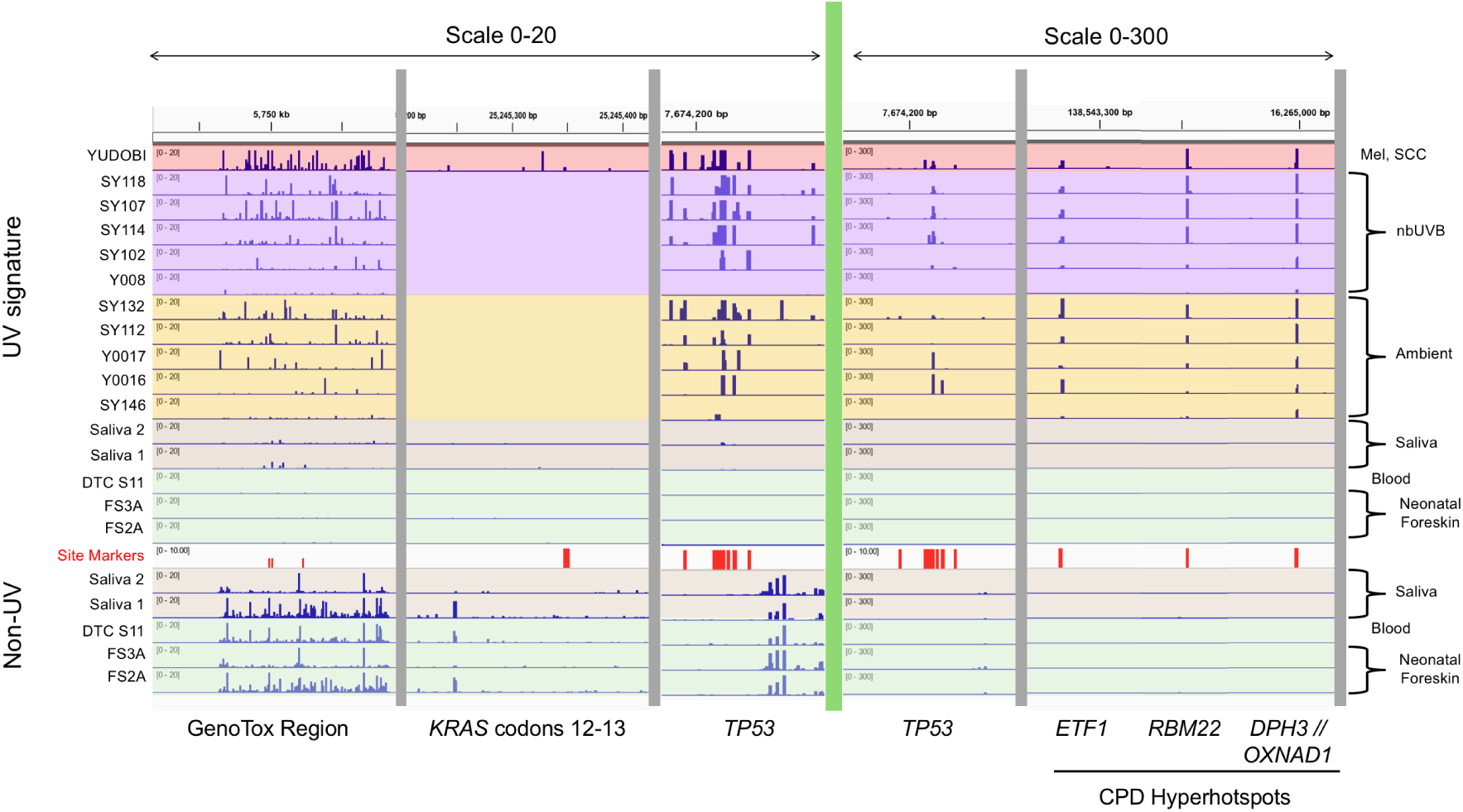
Mutation counts in four candidate genomic dosimeters: non-selectable region, oncogene, tumor suppressor gene, and CPD hyperhotspots. Integrated Genomics Viewer graphs show that non UV signature mutations (below red row) had a high background even in sun-shielded tissue, largely due to T→G and G→T substitutions. UV signature mutations (C →T at dipyrimidine sites) are shown above the red bars marking coordinates of tumor mutation hotspots, CPD hyperhotspots or, for GenoTox, elevated mutation frequency in some samples. GenoTox region: Commercially-optimized region for toxicology, putatively phenotype-free and found free of CPD hyperhotspots; C →T mutations in saliva are likely C deamination or G oxidation fortuitously at a dipyrimidine. Total mutation count increased with sun- or nbUVB-exposure. *KRAS* oncogene: Gain-of-function region codons 12-13. Mutation counts were low even in a patient with multiple skin cancers and were not localized to tumor mutation hotspots; oncogenes were dropped from the capture panel for later samples. *TP53*: This tumor suppressor gene, as well as *NOTCH1* and *NOTCH2*, had higher mutation counts than oncogenes especially at tumor mutation hotspots, likely due to clonal expansion; mutation count at individual sites increased with sun- or nbUVB-exposure. Codons 240 to 260 are shown. The same pattern is visible even at the 0-300 count scale (to right of green bar). CPD hyperhotspots: Shown are the promoters of *ETF1*, *RBM22*, and *DPH3*. CPD hyperhotspots were not mutated in sun-shielded tissue; in sun-exposed tissue they were typically mutated only at hyperhotspot nt, often more frequently than *TP53* sites. In some samples, the count extended beyond the 300 count scale shown. Y-axis, mutant DCS counts; note 15-fold larger scale to right of green bar. In these experiments, count 300 was a VAF of ∼1%. Abbreviations: DTC, commercial DNA Technical Control from blood buffy coat; Mel, melanoma; SCC, squamous cell carcinoma; nbUVB, therapeutic narrowband UVB exposure ranging from 5 months to 7 years; ambient, forearm exposed to ambient sunlight.

### CPD hyperhotspots are UV mutation hotspots in skin, often stronger than *TP53*

To investigate whether recurrent UV signature mutations arose from particular gene classes, notably cancer gene tumor hotspots or CPD hyperhotspots, we examined nt of interest within each class individually. For the cancer genes, the capture probes covered nt often mutated in skin tumors or sun-exposed skin, including nt found recurrently mutated in early phases of the present study. The CPD hyperhotspot probes were based on our earlier keratinocyte and melanocyte studies (*36, 37*). Successive experiments occasionally added or deleted probesets covering specific genes, so we restrict our comparisons of mutation classes to targets used in all studies.

To calibrate the magnitude of these VAFs against “null” sites that contain no CPD hyperhotspots and are presumed not susceptible to clonal expansion due to phenotypic selection, we used the entire GenoTox region as reference. No sites in this region are hotspots in tumors, so this calculation used all 845 dipyrimidine sites. (This large number also confers a lower detection limit, 3×10^-8^ per nt, than do the smaller panels of cancer gene or CPD hyperhotspot nt.) In **Fig. 3**, the left side of panels A and B shows each patient’s UV signature MF_maxI_ in the GenoTox region. For each target site, exposure to sun or nbUVB increases left to right between patient cohorts and within each cohort. The GenoTox region reached MF_maxI_ 10^-5^ per nt in sun-exposed skin; this is the upper end of the 10^-7^–10^-5^ per nt expected from classical tissue culture experiments using drug selection after a single UV exposure, although it would include any neutral drift *in vivo* (*32*). The typical VAF of an ordinary genomic site in sun-exposed skin is thus 500-fold lower than the 5×10^-3^ per nt detection limit of ordinary Illumina sequencing.

**Fig. 3.**
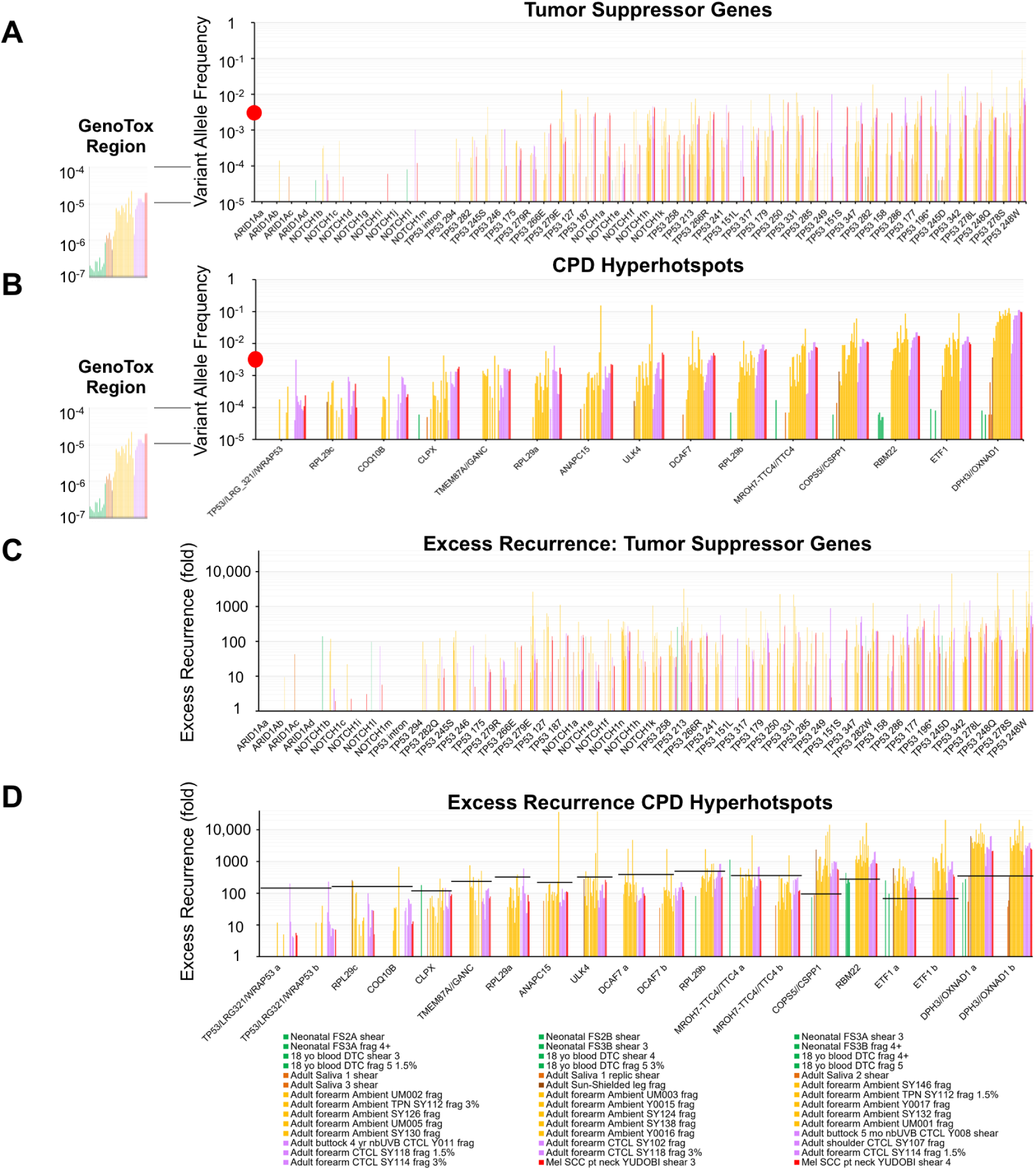
Variant Allele Frequencies for UV signature mutations across patients differ between tumor suppressor and CPD hyperhotspot genomic dosimeters. Each cluster pertains to a particular target nt; each vertical bar shows a patient’s value for that target nt. Patients are in order of increasing sun or nbUVB exposure or cancer status from left to right within each cohort and between cohorts (patient details in Fig. 4 and **Table S1**). Only targets used in all experiments are shown; more CPD hyperhotspots were added in later experiments. (A, B) Variant allele frequencies (VAF). (A) Tumor suppressor gene targets, chosen as tumor mutation hotspots plus sites reported to be frequently mutated in normal skin. They show a 10,000-fold range in UV-sensitivity. Sites were often unmutated in a biopsy; stochasticity is expected from clonal expansion of rare mutant cells. (B) CPD hyperhotspots. These also show a 10,000-fold range in UV-sensitivity, but most hyperhotspot mutations were present in nearly every sun-exposed biopsy and showed the same mutation pattern across exposure groups, with target sites differing only in scale. As a control, the far left cluster shows the UV signature mutant frequency per nt (MF) averaged across all cytosine-containing dipyrimidines in the reference GenoTox region, which lacks CPD hyperhotspots, features an ordinary UV mutation frequency, and is presumed to not confer a clonal expansion phenotype. Red dot indicates the detection limit of standard Illumina sequencing. Arrows indicate the VAF and MF detection limits for the amount of sample DNA and sequencing in these experiments, 7×10^-9^ per nt for the GenoTox region and 4×10^-5^ per nt for individual CPD hyperhotspot nt. Colors: green, neonatal foreskin, teenage blood; light brown, saliva; dark brown, sun-shielded leg; orange, ambient exposed forearm; violet, forearm from nbUVB-treated cutaneous T-cell lymphoma (CTCL) patients; red, melanoma and SCC patient. (C, D) Excess Recurrence of VAF relative to the ’ordinary’ and typically non-recurrent GenoTox region defined as having recurrence value 1.0. (C) Excess Recurrence for cancer hotspots; presence and magnitude are stochastic, with many sites absent in many samples. (D) CPD hyperhotspots. Values were not stochastic and the maximum value was typically close to the 100x-500x level expected from the enhancement of CPD frequency measured at that site in keratinocyte cultures exposed to nbUVB (horizontal bars). Yet VAFs at *ETF1*, *COPS5*, *RBM22*, and *DPH3* substantially exceeded the expected level, indicating that additional factors such as clonal expansion contribute. The greater detection sensitivity for Excess Recurrence reflects the fact that the GenoTox MF measurements are averaged over 845 Py-adjacent C, providing more sensitivity than single-nt VAF measurements at hotspot nt. Hence a VAF site in sun-shielded tissue having count 1 (non-recurrent) would need an Excess Recurrence of ∼845 before becoming detectable. For sun-exposed tissue, which is mutated more frequently in GenoTox, Excess Recurrence detection limits are lower, ∼100 for cohorts having brown bars and ∼10 for orange, violet, and red bars.

Oncogene gain-of-function mutations such as in *KRAS* or *BRAF* rarely appeared, nor did some genes seen mutated in skin in published studies, such as *FGFR3* and *FAT1*. Because an ultimate goal of the study is to measure individual UV exposures, oncogene targets were eventually omitted from the capture panel. The VAF of recurrent UV signature mutations at tumor mutation hotspots in the tumor suppressor genes *TP53* and *NOTCH1* reached 0.1–10% per nt (**Fig. 3A**), 100-10,000 fold greater than the 10^-5^ maximum of the GenoTox region. Because these genes did not appear in CPD hyperhotspot searches (*36, 37*), this excess reflects the clonal expansion of mutant cells having an altered phenotype rather than initial mutation induction. CPD hyperhotspot sites also showed mutations. Surprisingly, they also reached 0.1–10% per nt (**Fig. 3B**). Indeed, many CPD hyperhotspots had VAFs higher than those of the clones at *TP53* tumor hotspots. These and other mutation frequencies and ratios are collected in **Table S2**.

### Tumor suppressor gene mutations are sporadic across biopsies, but CPD hyperhotspot nucleotides are mutated in every sample

CPD hyperhotspots were mutated more consistently across patient samples than tumor suppressor genes were (**Fig. 3A, B**). This result identifies two classes of genomic regions that respond to UV exposure, which we will term “genomic dosimeters“:

1. Sporadic, where recurrent mutations rely on clonal expansion over time in those patient samples in which a rare founder mutation appeared. Clonal expansion is due to having a driver phenotype or being a passenger. For cancer genes, the sporadic driver behavior is expected.
2. Ubiquitous, where recurrence results from mutations independently induced by UV at high frequency in every patient. Any clonal expansion is superimposed on each of these frequent events, if the mutations are drivers, or on just a few of the events if expansion is as a passenger. Clonal expansion from neutral drift would also be ubiquitous, so an important reference standard is the mutation recurrence in a non-selectable genome region like GenoTox, where the MF would incorporate any neutral drift.

Ubiquitous dosimeters such as CPD hyperhotspots would directly measure accumulated UV dose. Measuring mutations at sporadic dosimeters such as *TP53* are less reflective of linear increases from dose and primarily measure the time required for exponential clonal expansion – either the duration of exposure or the time since exposure – as well as phenotypic selection for that mutation if it is itself the driver, or for another mutated driver if the scored mutation is a passenger. The distinction is important because many human and murine cancers depend on exposure dose to the 2nd or 3rd power (or less) but depend on exposure duration (e.g., cigarette pack-years) to the 5th or 6th power (*47–49*).

### The strongest CPD hyperhotspots exceed the VAF explainable by the site’s high CPD frequency

Assuming that the GenoTox region represents the mutation frequency typical of an ordinary dipyrimidine site and includes the extent of clonal expansion that results from neutral drift without selection, we used it as a standard to calculate the “Excess Recurrence” resulting from clonal expansion or from UV-hypersensitive nt (**Fig. 3C,D**). Tumor suppressor gene tumor hotspot nt exceeded the basal level by ∼20 fold on average, but across patients each hotspot nt ranged between 0-10,000 fold, i.e., they were often completely absent (**Fig. 3C**). This sporadic behavior fits with clonal expansion of rarely-induced driver mutations, and serves to calibrate the extent and variability of recurrence expected when mutant driver genes cause clonal expansion; a passenger mutation would be even less consistent. In contrast, each CPD hyperhotspot exhibited a consistent Excess Recurrence pattern varying mainly with exposure cohort and differing from other hyperhotspots by a scale factor (**Fig. 3D**). This uniformity is expected if the nt’s VAF was driven by CPD frequency rather than events selecting on the cell containing a mutant protein. The hyperhotspot Excess Recurrence was typically ∼100 fold but differed between hyperhotspots; some hyperhotspot target nt were 20-fold and some 10,000-fold.

A hyperhotspot’s maximum VAF value was typically close to the level expected from the enhanced CPD frequency measured at that site immediately after keratinocyte cultures were exposed to nbUVB (**Fig. 3D**, horizontal bars) (*37*). However, the elevated CPD sensitivity accounts for only the first 100-500 fold of the up to 10,000 fold excess of mutations at CPD hyperhotspots, leaving the higher-frequency hyperhotspots with a factor of up to 20-fold unaccounted for: *ETF1*, *COPS5*, *RBM22*, and *DPH3*. This surfeit could reflect poor repair, fast cytosine deamination, polymerase obstacles, or a clonal expansion phenotype conferred by the mutation. Which mechanism causes the excess VAF will become apparent later.

### Mutations in normal skin are pervasive even without sun exposure

Stochastic variation between cells in a biopsy is smoothed out by aggregating a large number of genome target sites. Mutation frequencies for the aggregated target regions were computed in two ways (*32, 50*). The minimum frequency of independently mutated genomes (MF_minI_) is defined as the frequency of *different* mutations, that is, the ratio: (number of different nt sites mutated, with alternative base changes at the same nt also counting as different)/(number of nt scanned). The number of bases scanned is the sum of coverage depth at each nt within the probe footprints (excluding captured flanking bases). In contrast, the maximum frequency of independently mutated genomes (MF_maxI_) is defined as the frequency of total mutations: (number of mutated nt)/(number of nt scanned). The number of mutated nt is given by the number of variant DCSs, each from one genome. If the mutant nt in a genomic target are each mutated just once, indicating stochastic rare events, then MF_maxI_ = MF_minI_. If MF_maxI_ > MF_minI_ , then some sites have recurrent mutations; either: a) the genome targets contain hotspots for DNA photoproducts or polymerase errors, leading to independently originating mutations at the same nt in different cells or b) mutant founder cells in the sample clonally expanded. In practice both events occur, so MF_minI_ is an underestimate of independent mutations and MF_maxI_ is an overestimate. Given the ∼14 kb total length of the target regions and the typical coverage depth of ∼25,000 DCSs per nt, the theoretical detection limit in these experiments (1/coverage) was an MF of 4×10^-9^ per nt. **Fig. 4** shows MFs averaged across all target genome regions that were used in every experiment (omitting targets deleted or added during development). It also expands the analysis beyond UV signature mutations.

**Fig. 4.**
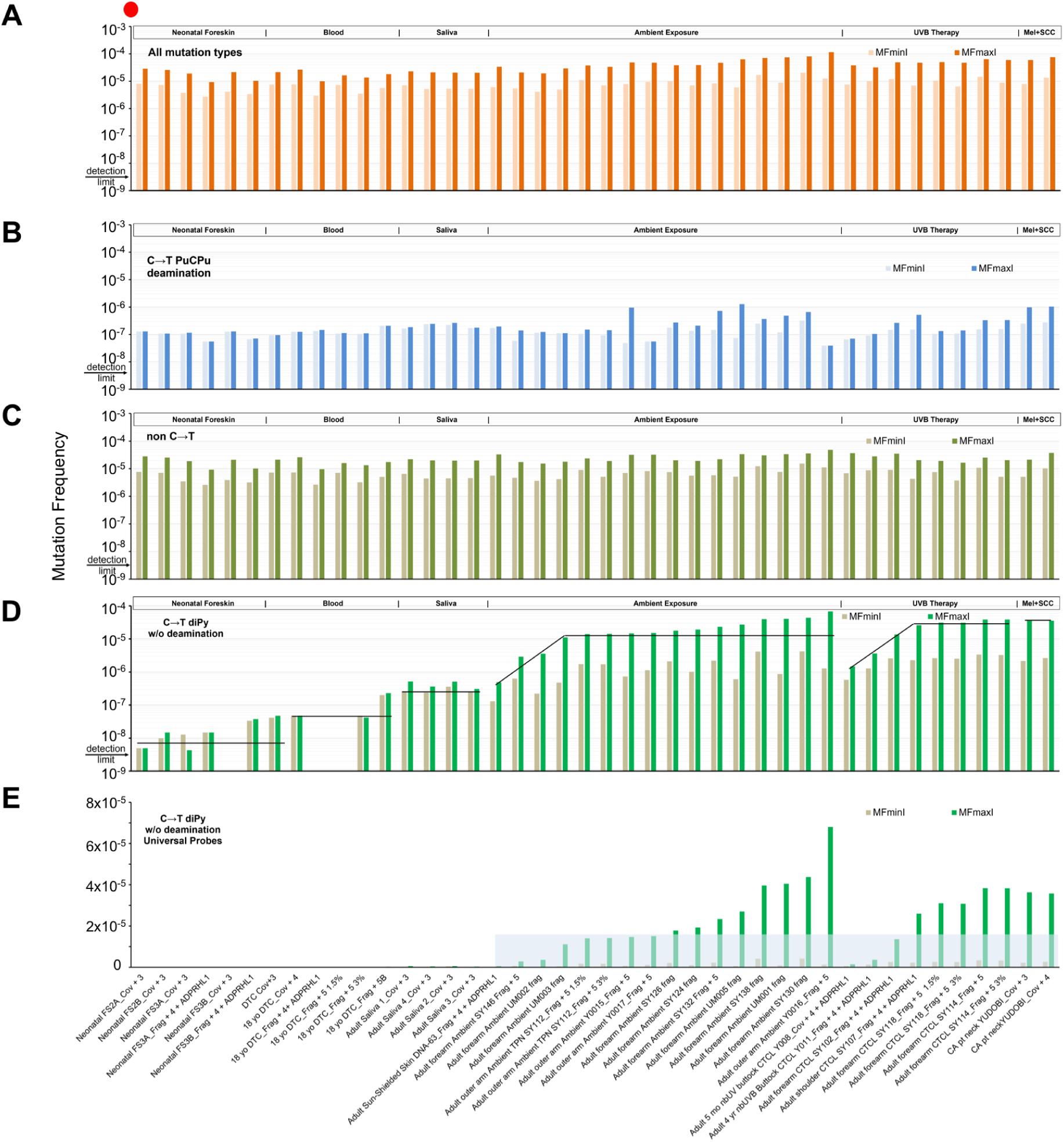
Recurrent UV signature mutations are a sensitive and specific indicator of sun exposure, yet are no more frequent than spontaneous mutations. For subjects having a range of sunlight or UV exposures, the minimum and maximum frequency of independent founder mutations per bp, MF_minI_ and MF_maxI_, were calculated across all captured target regions. (A) Mutation fraction for the aggregate of all mutation types, averaged over all targets used in every sample. Even MF_maxI_ is typically two orders of magnitude below the detection limit of standard next-generation sequencing (red dot). Both MF_minI_ and MF_maxI_ are substantial even in neonatal skin and sun-shielded body sites: the MF_minI_ of 3×10^-6^ per bp equals 18,000 different mutations per diploid cell. Aggregate MF_minI_ and MF_maxI_ varied little with UV exposure category. (B) Spontaneous cytosine deamination or G oxidation mutations (at purine-cytosine-purine sequences) occurred at a 100-fold lower frequency. They increased modestly with UV exposure in the nbUVB-treated CTCL group. (C) Non-C→T mutations, which include oxidatively induced mutations such as G→T and T→G, accounted for the first two logs of mutation frequency above cytosine deamination mutations or G oxidations leading to C→T; they showed even less variation with UV exposure than those. Sample repeats using shearing and fragmentase show fewer oxidative mutations with fragmentase, as intended. (D) UV signature mutations, arising from UV photoproducts joining adjacent pyrimidines, are quantified as C→T mutations at dipyrimidine sites after subtracting cytosine deaminations and G oxidations fortuitously occurring at dipyrimidine sites. Sun-shielded skin lacked recurrent UV mutations. On sun-exposed sites, the increase above unexposed skin was 30-500 fold for MF_minI_ and 100-8,000 fold for MF_maxI_. Mutations in blood and buccal epithelia/white cells at UV-like nt sites were elevated ∼5 and 20-fold relative to neonatal foreskin, respectively, for unknown reasons. (E) Plotting on ordinary axes shows the strong ability of UV signature mutations (C→T at dipyrimidine sites) to discriminate between sun-shielded and sun-exposed body sites. The mutation frequencies in ambiently exposed skin were less than in nbUVB-exposed patients, with the exception of an outlier who may be at risk. Abbreviations: FS, foreskin; DTC, commercial DNA technical control; nbUVB, narrow band UVB (311 nm); CTCL, cutaneous T-cell lymphoma; Mel+SCC, patient with melanoma and squamous cell carcinoma; shear, Covaris shearing; frag, fragmentase. Numbers refer to the successively refined target probeset panel used.

For the aggregate of all base substitution types, a logarithmic-scale histogram (**Fig. 4A**) shows that mutations were present at a frequency ≥3 logs above the detection limit in all samples, even in neonatal foreskin. These were typically recurrent (MF_maxI_ > MF_minI_), so they are not a stochastic assay background; that assay background is estimated to be ≤ 4×10^-10^ per nt (*51*). The MF_maxI_ of ∼1.5×10^-5^ per bp in neonatal foreskin equates to 90,000 different mutations per cell – even in a baby’s skin. MF_minI_, the independent mutations, were still half this, so neonatal mutations were not clonal expansions of rare mutations during embryo growth. These observations call attention to a remarkable degree of homeostasis in a tissue built of mutated genomes.

In sun-exposed skin, the MF_maxI_ of ∼4×10^-5^ or ∼240,000 mutations per cell in cancer-free individuals, is 10-fold higher than previously observed when detecting only clones having VAF > 1% (*3*). This frequency exceeds the upper limit of the 10^-7^-10^-5^ mutations per bp expected from a single UV exposure, based on classic drug-resistant colony assays (*32*).

Conversely, these mutation frequencies were 2-3 logarithms below the detection limit of standard next-generation sequencing, ∼0.5% per nt (red dot). Therefore, standard methods are restricted to very large mutant clones exceeding 0.5% at a particular nt. Importantly, without high fidelity methods even clones above 0.5% are nearly all spurious (*42*). With the more sensitive DuplexSeq method, it was clear that MF_minI_ and MF_maxI_ were surprisingly constant across UV exposure groups. To determine why, we next analyzed the mutations according to their physico-chemical origin.

### Cytosine deamination is modestly dependent on age and sun exposure

C→T mutations at non-dipyrimidine sites (purine-cytosine-purine sequences) originate from cytosine deamination at body temperature (*52*) or, less commonly, from guanine oxidation (*53*); their accumulation has been ascribed to age (*46*). **Fig. 4B** shows that these events constituted ∼1% of the total, with a slight increase between neonatal foreskin, adult saliva, and adult skin with ambient sunlight exposure in volunteers without skin cancer. They increased ∼3 fold with nbUVB treatment; this group also showed mutation recurrence. A UV increase at diPy sites would be expected because a CPD disrupts the 5-6 double bond and thereby accelerates deamination a million fold (*54, 55*). At non-diPy sites, an increase from UV is conceivable because, directly or via endogenous photosensitizers, UV oxidizes G and creates cytosine hydrates that disrupt the 5-6 double bond (*54, 56*). In sun-shielded skin, C deaminations or C→T mutations from G oxidation were non-recurrent (MF_maxI_ = MF_minI_), implying a random distribution without strong hotspots and, like the archetypal GenoTox region, without clonal expansion except in some ambiently exposed or nbUVB-treated patients.

### Non C→T mutations are nearly independent of sun exposure

Base substitutions other than C→T are typically due to spontaneous polymerase errors or oxidation-induced DNA damage, the latter causing mutations such as G→T and T→G (*57*). **Fig. 4C** shows that *non* C→T substitutions accounted for the two log increase above cytosine deamination in sun-shielded body sites. These polymerase errors and oxidative mutations increased little in skin exposed to ambient sunlight in volunteers without skin cancer. A pronounced sunlight increase might have been expected because oxidative mutations appear in laboratory UV-mutation studies (*57*) and because UV activates enzymes that synthesize reactive oxygen species (*31, 58*).

### UV signature mutations are specific for sun-exposed skin and reach high frequencies with normal sun exposure

To obtain the profile of true UV signature mutations, we subtracted from the “C→T diPy” profile the background C→T mutations unrelated to UV but arising because some C→T diPy mutations originating from cytosine deamination or G oxidation will have been only fortuitously located at dipyrimidines. These will be equal in number to C→T substitutions at PuCPu sequences (4 possible flanking sequences for each).

**Fig. 4D** shows that the MF of true C→T diPy mutations in never-exposed neonatal foreskin and presumptively-unexposed blood was ≤ 2×10^-8^ per nt and were non-recurrent (MF_maxI_ = MF_minI_); in some samples they were undetectable. Mutations in blood or in buccal epithelia and white cells at C→T diPy sites were elevated ∼5 and 20-fold relative to neonatal foreskin and blood, but still with little recurrence and thus random. These could be C deaminations or G oxidations uncompensated for by the correction factor above. In forearms exposed to ambient sunlight or nbUVB treatment, the MF_maxI_ of C→T diPy UV signature mutations increased 300-10,000 fold compared to neonatal foreskin. This induction compares to 2-3 fold observed when using standard NextGen sequencing containing spurious mutations arising from damage in the DNA template (*43, 59*). This histogram also makes it clear that replicate Duplex Sequencing samples were reproducible even when different probesets or DNA fragmentation methods were used. The MF_maxI_ reached 3×10^-5^ per nt, or 30 per Mb and 180,000 per cell, presumably reflecting multiple exposures over a lifetime. Assuming 100 exposures – which is plausible for a minimal erythemal dose – and no loss of mutant cells, this value extrapolates to an MF of 1×10^-7^ for a single outdoor exposure. This is at the lower end of our estimate from the literature, which corresponded to non-lethal UV doses.

The data reveals a limitation to the use of MF_minI_ at high exposures. When all distinct genomic coordinates that are mutable have been mutated at least once, MF_minI_ = (genome coordinates mutated)/(genome coordinates scanned * coverage per nt) will equal 1/(coverage per nt) and thus a plateau in MF_minI_ will result. For 25,000x coverage, this value is 4 × 10^-5^ and is in fact the maximum MF_minI_ observed in C→T diPy mutations in **Fig. 4D**. In human skin, this plateau regime in which all available genomic coordinates were mutated at least once was reached even in ambient-exposure patients. A side effect is that MF_minI_ has limited usefulness as a metric for independent mutations induced by UV exposure, so better metric is the Excess Recurrence used above.

Plotting on ordinary axes (**Fig. 4E**) highlights the UV-exposure specificity of the C→T diPy mutations. It also indicates a provisional “normal range”, with most nbUVB-treated patients falling above this range. One outlier ambient-exposure volunteer may be a candidate for at-risk monitoring; a larger study is needed in this regard.

### Mutation creation and clonal expansion are both concentrated at specific nucleotides

#### UV mutation creation was localized

The MF_minI_ for inducing independent UV mutations across all universal target regions in sun-exposed skin was 0.2×10^-5^ (**Fig. 4**, **Table S2**), the same as the MF_minI_ in the putatively non-selectable, CPD hyperhotspot-free GenoTox region and within a factor of two of that in cancer genes and the 120 nt long regions containing CPD hyperhotspots (**Table S2**). This equivalence implies that most mutant nt in our target regions, including cancer genes, resemble GenoTox in not being selected and not containing CPD hyperhotspots.

#### Recurrence due to neutral drift was undetectable

Neutral drift would affect every cell in a tissue sample, so even non-selectable mutations would be recurrent. In the non-UV-sensitive, putatively non-selectable GenoTox region, the total MF_maxI_ in sun-exposed skin reached 0.5×10^-5^ per nt (**Table S2**). An alternative to MF_maxI_, the average VAF across the GenoTox region, was similar, 10^-5^ (**Fig. 3**). Recurrence, quantified as the ratio MF_maxI_/MF_minI_, was thus 2.5-5.0 fold, giving a Poisson average recurrence of 1.5-4.0 fold. At Poisson average recurrence 3.5 and a typical coverage of 25,000, recurrence would be statistically significant at p<0.05 for an initial mutation count giving VAF above 3.2×10^-4^. Only 2 of the 2401 nt in the GenoTox region exceeded that value in more than one biopsy (two biopsies), so the observed ∼3.5-fold average recurrence does not represent a statistically significant neutral drift. Neutral drift in the GenoTox region was thus undetectable. Because neutral drift is a property of a tissue, rather than a particular mutation, this observation means that any statistically significant mutation recurrence in another gene target relied on events other than neutral drift.

Estimating an upper limit to neutral drift takes into account the fact that measured mutation recurrence does not fully measure the extent of clonal expansion. In our experiments, the ∼25,000 genomes underlying the coverage constituted 1/8 of the ∼200,000 genomes in the original 0.5 ug DNA biopsy. Sequencing 8-fold more genomes would, at MF_minI_ 0.2×10^-5^, be unlikely to produce a second, independent founder mutant at a particular nt. However, it would provide an 8-fold larger sampling of the original founder’s clonal progeny. The degree of clonal expansion due to neutral drift was therefore 8-fold larger than measured. If we conservatively assume that every count 1 mutation was an undersampled neutral drift clone, neutral drift was at most 8 fold. This is less than the 32-fold seen in the course of a year for tagged cells in mouse skin, where it reaches a time-independent equilibrium (*60*). By referring other recurrence measurements to the rate in the GenoTox region (Excess Recurrence), we measure recurrence independent of any contribution from neutral drift.

#### Recurrence in cancer genes is modest

Similar calculations for the entire probe region covering tumor mutation hotspot nt yielded average Recurrences of 10-fold (**Table S2**) and thus average clonal expansion of 80-fold.

#### Recurrence near CPD hyperhotspots is modest

The entire probe region covering CPD hyperhotspot nt yielded average Recurrences of 20-fold (**Table S2**). For recurrence arising from independently-arising mutations, the above effect of sample size does not apply. Therefore, the previously mentioned 10,000-fold Excess Recurrences of particular nt with respect to the GenoTox control region is specific to those nt, the tumor hotspots or CPD hyperhotspots. Is the excess recurrence of either hotspot class explainable as simply being a passenger mutation?

### Passenger clones are vanishingly rare in skin

If a mutation has no clonal expansion phenotype itself, it can clonally expand beyond neutral drift by being a passenger. UV-induced mutations at CC sequences provide an internally-controlled test for phenotype vs passenger, applicable *in vivo*. Passenger test: At a CC site in which C_2_→T is a synonymous mutation, occurrence of both C_1_→T and C_1_C_2_→TT (in different genomes in the sample) implies that: C_1_→T has a selectable phenotype, biochemistry can make C_2_→T mutations at this site; and C_2_→T can be expanded as a “*cis* passenger” by selecting on the C_1_ mutation’s phenotype. Crucially, any synonymous C_2_→T mutations that recur singly rather than as part of the CC→TT recur because the silent mutation was a passenger in a cell mutated in some other gene that drove clonal expansion. The frequency of singlet C_2_→T recurrences at such test sites (occurrences – 1) gives the frequency of conventional “*trans* passenger” mutations. (And *mutatis mutandis* if the synonymous mutation is at C_1_.)

Two sites of this type are present in *TP53* tumor hotspot codons 248 and 247 (in reference genome order). The DNA sequence is 5’ cCG**|**g 3’, where lower-case indicates the third position of a codon where its C→T mutation is silent (**Table 1**). The passenger mutation frequency in these biopsies was 0. Sites of this type are also present in *TP53* tumor hotspot codons 177 and 342. The silent C→T mutation was observed singly only once and was non-recurrent. Because these *TP53* missense mutations drive clonal expansion (*2*), the mutation counts in the full biopsy were, as discussed above, 8-fold the numbers shown in the Table.

**Table 1.** Consistent absence of recurrent passenger mutations at silent C→T substitutions in *TP53* in sun-exposed skin. Codons 247-248, 177, and 342. Mutated base(s) in bold; silent mutations in lower case; vertical bar is the codon boundary. Samples arranged in order of increasing patient exposure to sun or nbUVB. DNA sequence is given as the reference genome top strand; the TP53 protein is coded on the bottom strand, so codon order is right to left. hg38 coordinates chr17:7,674,219-222; 7,675,081-082; and 7,670,685-686.

| Mutation | Amino Acid Change | Y0017 | SY132 | Y0016 | SY102 | SY107 | SY118 | SY114 |
| --- | --- | --- | --- | --- | --- | --- | --- | --- |
| c**CG** \| g → c**TG**<br>\| g | R248Q | 0 | 18 | 27 | 52 | 22 | 24 | 30 |
| c**CG** \| g → t**TG**<br>\| g | R248Q | 0 | 5 | 0 | 12 | 28 | 8 | 2 |
| c**CG** \| g → t**CG**<br>\| g | synonymous (248) | 0 | 0 | 0 | 0 | 0 | 0 | 0 |
| c**CG** \| g → c**CA**<br>\| g | R248W | 246 | 26 | 1017 | 13 | 107 | 54 | 69 |
| c**CG** \| **g** →<br>c**CA** \| **a** | R248W | 0 | 35 | 9 | 0 | 92 | 68 | 127 |
| c**CG** \| **g** →<br>c**CG** \| **a** | synonymous (247) | 0 | 0 | 0 | 0 | 0 | 0 | 0 |
| g**G** → g**A** | P177L | 10 | 20 | 0 | 3 | 44 | 57 | 30 |
| g**G** → a**A** | P177L | 3 | 6 | 0 | 2 | 44 | 11 | 0 |
| g**G** → a**G** | synonymous | 0 | 0 | 0 | 0 | 0 | 0 | 0 |
| **C** \| c → **T** \| c | R342stop | 6 | 8 | 0 | 0 | 66 | 7 | 177 |
| **C** \| c → **T** \| c | R342stop | 0 | 0 | 0 | 0 | 12 | 34 | 32 |
| **C** \| c → **C** \| t | synonymous (341) | 0 | 0 | 0 | 0 | 0 | 0 | 1 |

We conclude that for sites mutated at typical frequencies – including *TP53* tumor hotspots – non-selectable mutations are generated so infrequently in biopsies of this size that clonal expansion by being a passenger is orders of magnitude rarer than direct clonal expansion. Correspondingly, a site often recurrently mutated above the neutral drift level is likely to have a phenotype enhancing cell survival, mutation creation at a CPD, or clonal expansion.

This lack of passengers agrees with the 0.3×10^-5^ per nt frequency of inducing independent UV mutations at ordinary sites: Given a UV MF_minI_ of 0.6×10^-5^ per cell at a particular nt, and assuming that any of 1000 nt in one cancer gene or another can lead to clonal expansion if mutated, the frequency of the original passenger event for a particular mutation would be (0.6×10^-5^)*(1000*0.6×10^-5^) = 4×10^-8^ per cell; this is 1 per 100 cm^2^ of skin (*61*). A CPD hyperhotspot with 100-fold UV sensitivity would be a passenger on a rare driver once per cm^2^ of skin. In comparison, a biopsy yielding 600 ng of DNA came from only 0.02 cm^2^ containing 100,000 cells.

### CPD hyperhotspot mutations have a clonal expansion phenotype *in vivo*: allele imbalance

DNA sequence reads from homogenized tissue do not readily reveal whether an individual recurrent mutation arose from independent events or a rare mutant cell that clonally expanded. We used two strategies to identify clones of mutants at individual nt, both using the fact that independent mutational events are stochastic.

First, independent stochastic events will be distributed equally between alleles. In contrast, a clone derived from one founder will create an asymmetry, especially when the number of founders is not large. Statistically significant allele asymmetry indicates clonality. Ambiently-exposed non-patients and nbUVB-treated patients occasionally carried SNPs in the regions studied here; curiously, skin tumor mutation hotspot regions were devoid of SNPs. Allele counts and p values for allelic imbalance at sites known to be mutable (i.e. having at least 1 mutation count in one patient) and having a total count ≥3 are presented in **Table S3**. These criteria were met by 97 SNP-adjacent sites. Typically, imbalance was statistically significant when the total count was ≥6 and at least one allele had count ≥5.

Allele imbalance was frequent. As expected, 70% of the 13 qualifying mutations in *TP53* showed statistically significant allele imbalance, reflecting its role as a cancer gene that drives clonal expansion, particularly in the presence of chronic UV exposures (*21, 23, 25*). Surprisingly, allele imbalance was statistically significant in 41% of the 81 mutations in CPD hyperhotspot regions, despite needing to overcome the law of large numbers in these highly mutable sites. Moreover, the same imbalanced CPD hyperhotspot sites appeared in multiple patients. The prevalence of clonal expansion suggests that the CPD hyperhotspot mutation itself has a clonal expansion phenotype. These imbalanced mutations also included nt that are not CPD hyperhotspots, but were nearby, suggesting that the entire region of the RNA processing gene promoters resembles *TP53* in having a clonal expansion phenotype when mutated. The extent of clonal expansion is indicated by the allele’s mutation count if one founder is assumed; this is a reasonable approximation when the other allele’s count is 0 or 1. This factor is then multiplied by the sampling factor of 8x discussed above to estimate the actual extent of clonal expansion in the tissue.

**Fig. 5A** plots the p value for allelic imbalance as a function of total mutation count. Several points are evident:

- Allelic imbalance increased with mutation count, as it would from large clones, rather than decreasing as the law of large numbers would dictate if mutations were independent stochastic events with no clonal expansion.
- *TP53* mutations were typically monoclones, mutated on only one allele, as expected if a rare event clonally expanded.
- CPD hyperhotspots were occasionally monoclones, often near-monoclones, as well as occasionally allele-balanced. This is the behavior expected from initial mutation frequency being higher than in *TP53* and if occasional clonal expansion was superimposed on the creation of independent mutations.
- The CPD hyperhotspot used in all experiments that had the most statistically significant allele imbalance (p=10^-12^) was *ETF1*. For its two hyperhotspot nt, 92% (48 of 52) of the sample-hyperhotspot combinations examined in sun-exposed skin acquired more UV signature mutations than accounted for by the sites’ 90-fold elevated CPD formation (**Fig. 3**). The excess recurrence was as great as 20,000 fold; the median was 430 fold, a 5-fold increase over expected. Again correcting for the 1/8x sampling factor, it is most likely that 40-fold was the typical clonal expansion of individual founder mutations at this CPD hyperhotspot.

**Fig. 5.**
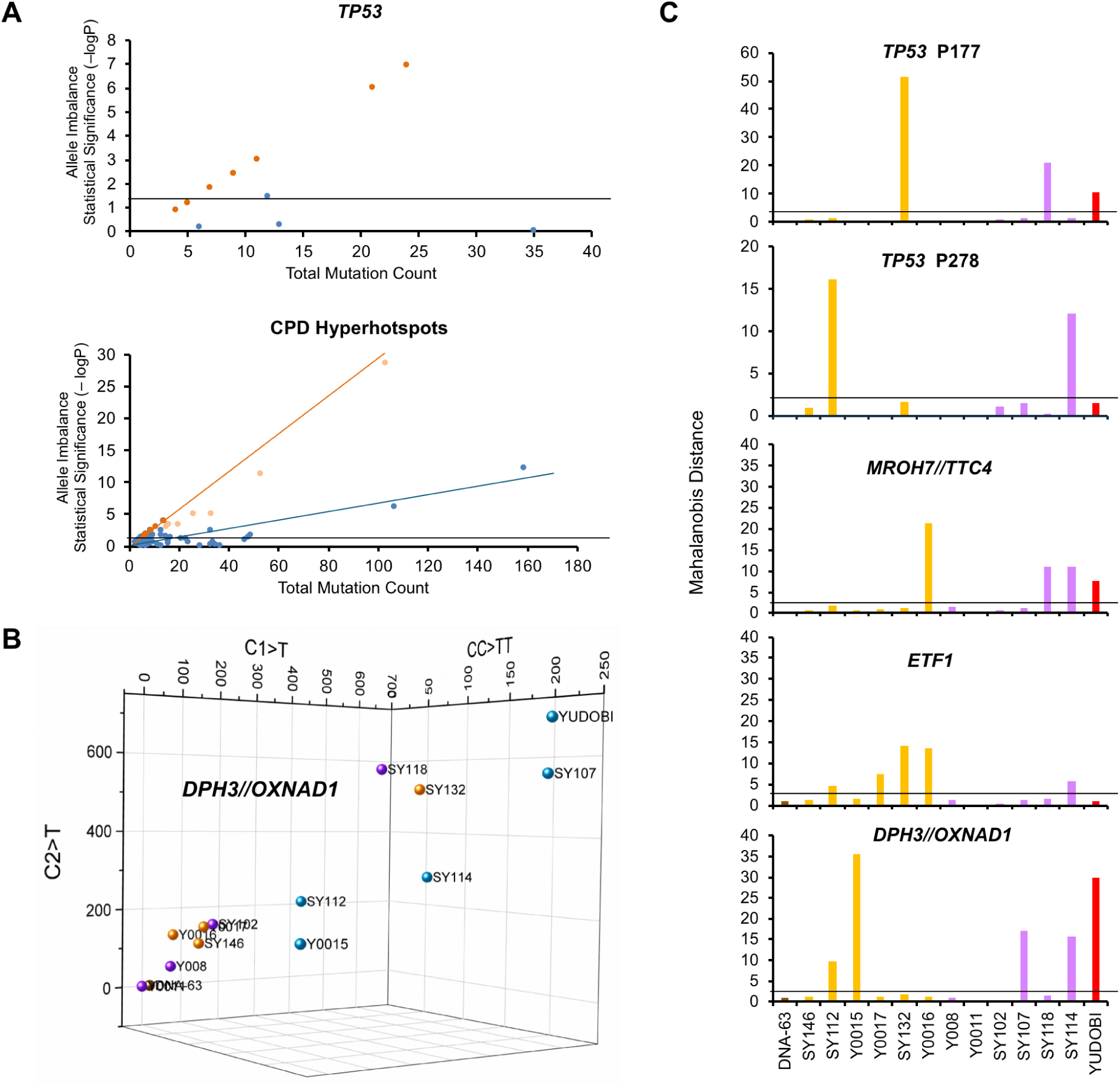
Non-stochastic mutation distributions due to clonal expansion. (A) Allele imbalance. Mutations in normal sun-exposed skin were often unequally distributed between two alleles identifiable by a SNP, indicating they arose from non-independent events such as clonal expansion of a founder mutant. Dark gold circles, monoclones having 0 mutations on the other allele; light gold, near-monoclones; gray horizontal line, p = 0.05. Even CPD hyperhotspot mutations exhibit clonal expansion behavior. (B) Inter-patient imbalance. At a CC sequence in the *DPH3* CPD hyperhotspot, the count of UV-induced 5’C→T, CC→TT, and 3’C→T mutations in sun-exposed skin increased with patient sun or UV exposure; their ratio typically did not change, resulting in a line. Outlier samples departed significantly from the predominant ratio of the three mutation types at that site (light blue spheres), indicating a non-stochastic event such as clonal expansion. 2D projection of 3D Origin plot of data from the CPD hyperhotspot in *DPH3*. Brown, sun-shielded leg; orange, ambient forearm exposure; violet, nbUVB-treated CTCL patients; outliers Y0015 and SY112 were ambient, SY114 and 107 were nbUVB, and YUDOBI was uninvolved skin of a patient with melanoma and squamous cell carcinoma. (C) Mahalanobis distance. Divergence of the sample’s mutation count from the centroid of data. Values above the blue horizontal line at 3.1 are statistically significant. The outlier sites varied from sample to sample, indicating that the magnitude of clonal expansion varied, even in *TP53,* and was not a biochemical property of the site or a propensity of the patient.

### CPD hyperhotspot mutations have a clonal expansion phenotype *in vivo*: patient imbalance

The second strategy for identifying clonal expansion, one not requiring SNPs, compares patients rather than alleles. A CC site has three options for UV-induced mutations: 5’ C→T, CC→TT, and 3’ C→T. Their ratio might vary between CC locations due to biochemistry, but at any particular CC stochastic new mutations will occur at the same ratio in each patient. Inter-patient disparity indicates a non-stochastic event such as clonal expansion. CC sequences were common at tumor mutation hotspots of *TP53* and *NOTCH1* and at CPD hyperhotspots. Ratios were compared using the Mahalanobis distance from the centroid of the data (Methods).

For *TP53* tumor hotspots, the algorithm was unable to compute a centroid of the data for most sites, indicating that either many patient samples had 0 counts or detectable counts relied on being clonal and thus nearly all counts were outliers. However, codons 177 and 278 did allow this computation and 3 of 10 and 2 of 9 non-zero samples, respectively, were statistically significant outliers. In comparison, all CPD hyperhotspots examined produced outliers (**Fig. 5B**), indicating that significant clonal expansion was superimposed on the high mutation rate stemming from their 100x CPD frequency. Which hyperhotspots were outliers varied from sample to sample (**Fig. 5C**), indicating that the magnitude of a founder’s clonal expansion varied, even in *TP53,* and that it was not a biochemical property of the site. The rare driver mutations at *TP53* tumor hotspots showed extensive clonal expansion in only about half the samples in which they appeared at all (**Fig. 5C**), indicating that either the mutation was recent, additional events are required, or expansion was followed by regression (*21, 25, 60*).

### In cell culture, mutation hotspots arise at CPD hyperhotspots days after a single low-dose UVB exposure and some increase exponentially with time

To assess whether the high UV signature MFs we observed required the patients’ years of low-dose sun exposure, we irradiated a 30% confluent culture of near-primary human neonatal keratinocytes with known UVB doses up to 8 times over 4 weeks. To avoid an atypical result, we used a commercial mixed-donor culture consisting of at least 3 donors, evidenced by SNPs in the Day 1 sample falling into one of three frequency categories.

Most CPD hyperhotspots developed mutations with VAF up to 10^-3^ after one exposure, with several continuing to increase with continued chronic irradiation. Other CPD hyperhotspot sites required four exposures over two weeks (**Fig. 6A**). VAF reached as high as 3% in four weeks. The earliest-developing sites included *ETF1*, *COPS5*, and *DPH3*, the sites whose mutation Excess Recurrence exceeded the level accounted for by the CPD hypersensitivity (**Fig. 3D**). During the third week, the VAF at some sites decreased or disappeared. This decline did not reflect an effect of the mutation, because the timing and extent of these decreases corresponded to the timing and extent of decrease of two SNP families (**Fig. S2**). This selection evidently reflected the donor’s growth rate or plating efficiency, rather than UV sensitivity, because it also occurred in the unirradiated controls. Both the emergence of frequent mutations and the disappearance of some mutations were greater at the higher dose. These results experimentally confirm the model of high-frequency independent mutations at ETS-1 binding sites that we and others (*36, 41, 62*) have proposed.

**Fig. 6.**
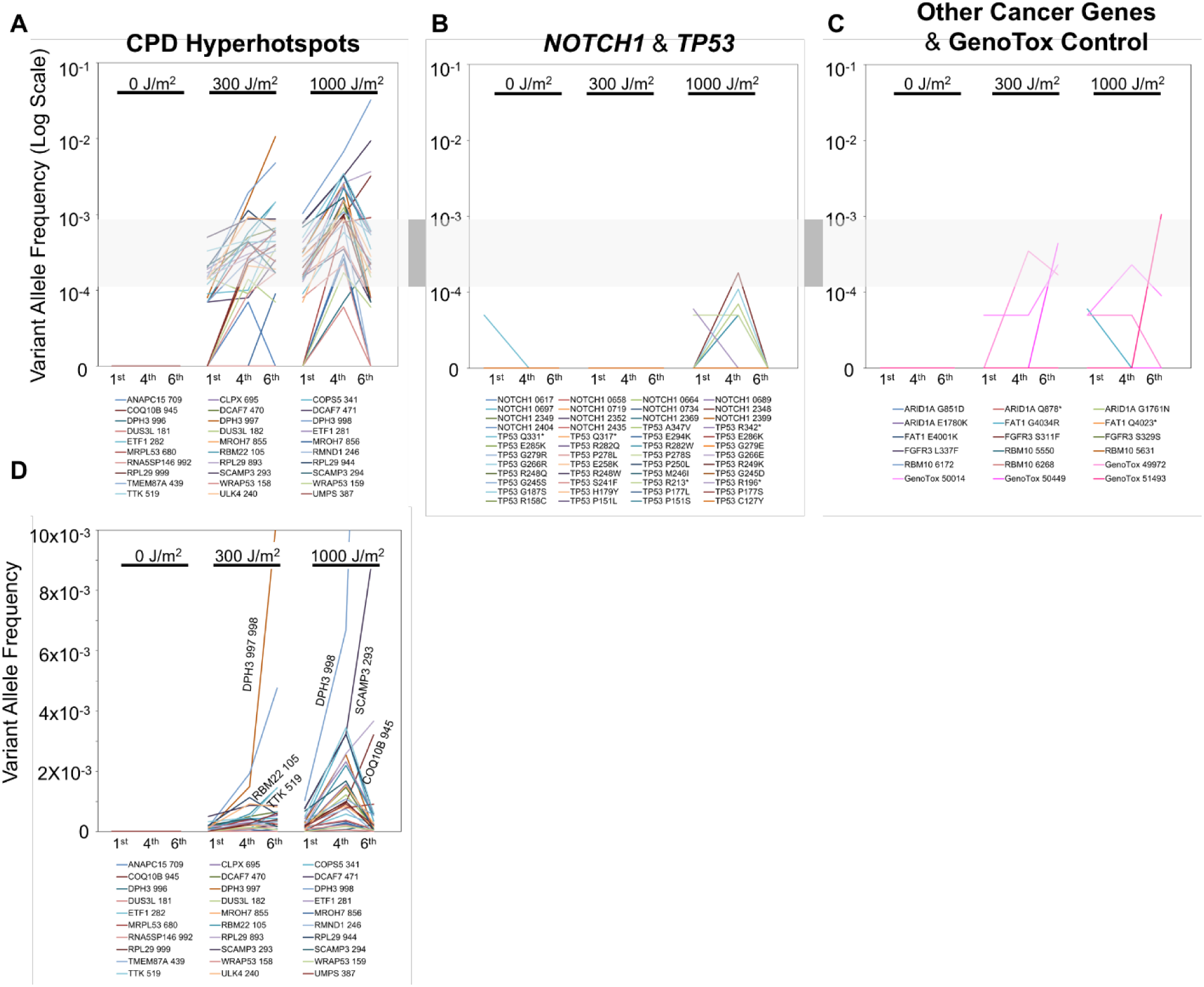
Prompt and exponential emergence of UV signature mutations at CPD hyperhotspots. Keratinocytes chronically irradiated with UVB developed mutations at CPD hyperhotspots after one irradiation; the highest VAF sites increased exponentially thereafter, indicating clonal expansion rather than cumulative independent mutations. Mutations in oncogenes, tumor suppressor gene tumor hotspots, and a phenotype-less control region were much less frequent and appeared much later, indicating that the CPD hyperhotspots were not their passengers. Log-scale plot of VAF for : (A) CPD hyperhotspots, (B) *NOTCH1* and *TP53* tumor suppressor genes, and (C) oncogenes, additional tumor suppressor genes, and the phenotype-less GenoTox control region. (D) Non-log plot showing exponential VAF increases for CPD hyperhotspots.

The strongest CPD hyperhotspots increased log-linearly (**Fig. 6A**), implying exponential growth rather than a linear increase with time and additional exposure. The magnitude of this non-linearity is evident on a non-log plot (**Fig. 6D**). In skin, the Excess Recurrence of the strongest CPD hyperhotspots (**Fig. 3D**) could have been attributed to the site’s poor repair, fast cytosine deamination, or being an obstacle to polymerase. The exponential increase observed across multiple timepoints *in vitro* rules out these mechanisms and leaves clonal expansion. Therefore, although CPD hyperhotspot mutations could be passengers by virtue of arising frequently, they do not need to be.

### Mutations at tumor mutation hotspots and phenotype-free regions arise too late and too rarely to be drivers for CPD hyperhotspot mutation passengers

A priori, this clonal expansion could be as a passenger or due to a driver phenotype conferred by the mutation. The passenger mechanism was shown above to be numerically unlikely at ordinary mutation frequencies, but would be possible for a CPD hyperhotspot if a 500-cell driver clone had arisen in advance in every sample, and thus in every sample-sized skin region in the donor. Strikingly, *in vitro* mutations in tumor suppressor genes and oncogenes arose two weeks later than at CPD hyperhotspots, if at all, and the maximum VAFs at tumor suppressor genes were 1000-fold lower than at CPD hyperhotspots (**Fig. 6 C, D**). Indeed, their behavior was not distinctively different from that of the putatively phenotype-free GenoTox region (**Fig. 6D**). Therefore, CPD hyperhotspot mutations cannot have been passengers on these cancer genes. It is also unlikely that they were passengers on any other cancer genes, inasmuch as no cancer genes appeared in the list of high CPD frequency sites in keratinocytes, melanocytes, or fibroblasts (*36, 37*). Evidently, clonal expansion of rare mutations is significantly slower than accumulation of recurrent mutations resulting from recurrent CPDs at CPD hyperhotspots and require more than 30 days and 6 chronic exposures to become appreciable. Independent CPD hyperhotspot mutations can arise quickly in sun-exposed keratinocytes, do not require years of sun exposure, and can become more pervasive in sun-exposed skin than cancer gene mutations. The rapidity with which these mutations arose, and the fact that some sites had higher MF than accounted for by the CPD sensitivity (**Fig. 3D**), imply that CPD hyperhotspot mutations in RNA processing gene promoters have a mini-driver phenotype *in vivo* (*63*).

## Discussion

Quantifying the mutational events in normal skin illuminates the onset of tumor development and identifies genomic dosimeters – regions of DNA that quantify an individual’s past sunlight exposure and ostensible risk of future skin cancer. Surprising observations were the lack of passenger mutations or neutral drift clonal expansion, high mutation frequency in neonatal skin, a frequency of UV signature mutations at CPD hyperhotspots exceeding that at tumor mutation hotspots in tumor suppressor genes and appearing consistently in every biopsy, and a clonal expansion phenotype of CPD hyperhotspot mutations.

### Genomic dosimeters

Considering MF_maxI_, the metric that includes recurrent mutations at the same nt, as a genomic dosimeter, we find that neonatal skin already has a level of 1.5×10^-5^ per nt when all base-change types are included, or 90,000 per cell. This value may be underestimated inasmuch as our scale of ∼500 ng DNA and ∼200M sequencing reads detects ∼1/8 of the single cells that mutated but didn’t clonally expand. UV signature mutations, C→T at a dipyrimidine, were ≤ 1/4000 of this value, perhaps due to other causes but fortuitously located at this motif, and they rose 8000-fold to 3×10^-5^ in skin exposed to ambient sunlight over decades. For most of the ambiently exposed volunteers, intense sun exposure was primarily in youth, so UV signature mutations are a persistent, sensitive, and specific, genomic dosimeter.

Yet this MF_maxI_ value was only 2-3 fold higher than the total of mutations present in neonatal skin, raising twin questions: Why are UV signature mutations the ones primarily associated with skin tumors? How does skin function as well as it does when it is a sheet of mutant cells? The former fact may indicate that UV exposure also drives clonal expansion of the UV-mutated cell (*21–25*), whereas spontaneous oxidative damage or polymerase errors do not. Homeostasis despite huge numbers of errors indicates that life involves a level of robustness that we do not yet understand.

Focusing on initial mutation creation, the average MF_minI_ for independent UV signature mutations was ∼0.3×10^-5^ per nt whether in the phenotype-less GenoTox region, tumor suppressor genes, or promoter regions near CPD hyperhotspots. The regions diverged only for MF_maxI_, due to recurrence of the same mutation in multiple cells. This recurrence was due to clonal expansion for tumor suppressor gene mutations, but for CPD hyperhotspot regions it was due to high frequency independent production of CPDs at the same nt in separate cells.

At specific nt, these recurrences reached 0.1-10%. A striking dichotomy was apparent between genome regions. Tumor mutation hotspot nt in tumor suppressor genes reached this level via 30-3,000 fold clonal expansion, but only in some biopsies, making them “sporadic dosimeters”. In an assay, many of these would need to be examined in any particular biopsy; their mutation frequency would reflect a low creation frequency and a long time needed for clonal expansion, so the net MF measures primarily time. Surprisingly, CPD hyperhotspots reached the same mutation frequencies or higher, and acted as “ubiquitous dosimeters” that gave a signal in every biopsy; these were reflective of initial dose.

### Mini-drivers

Regarding the driver of clonal expansion, the common expectation is that a mutation is either a cancer driver (setting aside for the moment whether a cancer driver is always an expansion driver), a passenger, or a beneficiary of neutral drift affecting all cells in a region of skin. The phenotype-free GenoTox region showed that neutral drift was undetectable. CC sites at which one of the possible C→T mutations was silent showed that passenger mutations almost never occurred, even in *TP53* (the silent C→T mutation was observed only in CC→TT). These facts leave clonal expansion phenotypes as the explanation for genes like *TP53*. For CPD hyperhotspots, the null hypothesis would be that they have no phenotype and they attain their high MF as passengers (*62*) or due to their high CPD frequency (*36, 37*). Yet even the CPD hyperhotspot mutations possessed a clonal expansion phenotype, revealed in the context of non-stochastic imbalance between alleles or between patients. That phenotype was also evinced by the rapid and exponential growth of CPD hyperhotspot mutations in culture. CPD hyperhotspot mutations thus resemble one type of mini-driver (*63*).

Can mini-driver mutations at CPD hyperhotspots facilitate cancer? Given the 0.3×10^-5^ per nt frequency of inducing independent UV mutations at ordinary sites (and thus a UV MF_minI_ of 0.6×10^-5^ per cell at a particular nt), and the epidemiologic evidence that many cancers involve ∼6 carcinogen-related mutational steps, the frequency of the same cell acquiring two particular mutations is (0.6×10^-5^)^6^ = 5×10^-32^, if these are independent. If any of 1000 nt in one cancer gene or another can be mutated for each step, it is 5×10^-14^. But a person’s 0.5 m^2^ of sun-exposed epithelium contains only 2.5×10^10^ cells (*61*) and U.S. skin cancer incidence is 20% not 10^-3^.

Clonal expansion evades this rarity constraint and enables carcinogenesis (*64–69*). If each cancer gene’s mutation led to 10x clonal expansion that increased the likelihood that one of the daughters receives a second cancer gene mutation, the discrepancy between 20% and 10^-3^ would quickly be overcome. However not all cancer genes autonomously drive clonal expansion, the *RAS* family being an example, and even mutant *TP53*’s clonal expansion requires non-mutational effects of UV exposure (*21, 25*) and may potentially need to occur in a specific order. In contrast, if any of 200 100x-mutable CPD hyperhotspots can be mutated to drive clonal expansion, and that expansion is 10-fold, the probability of creating a mini-driver is (200*100*0.6×10^-5^) and the probability of acquiring each successive mutant cancer gene in the same cell is (10*1000*0.6×10^-5^), for a net of 6×10^-9^ for the six mutations. There are then 150 skin cancer cells per person. Pervasive mini-drivers modestly driving clonal expansion can let a single cell acquire multiple UV mutations.

For mini-drivers involved in cell housekeeping functions, as CPD hyperhotspots in RNA processing genes are, it is plausible that they can synergize or at least sum. Because of the CPD hyperhotspots’ high mutation frequency, co-occurrence is likely. An estimate of the likelihood that three of 200 100x-mutable CPD hyperhotspot nt are mutated in the same cell by the same UV exposure is (200*100*0.6×10^-5^)^3^ = 2×10^-3^ per cell, 9000 per cm^2^, or 200 cells per 600 ng sequencing sample. We conclude that UV mutagenesis targets mini-drivers that modestly but pervasively drive clonal expansion *in vivo*.

### Potential implications and limitations of these results

The pervasive presence of mini-driver mutations that favor clonal expansion means that random absorption of UV photons across the genome nevertheless results in a non-random emergence of mutations. The clonal expansion facilitated by these mutations means that even oncogenes and tumor suppressor genes that do not themselves cause clonal expansion can have their rare initial mutation frequency amplified to a substantial mutation burden in the tissue, thereby increasing the likelihood of acquiring the other mutations or epigenetic events required for cancer. The sensitivity of these measurements means that UV-induced mutations in skin can serve as an objective measure of an individual’s sun exposure history. Limitations to these results are that mutation frequencies at an instant may reflect an equilibrium between creation and elimination of mutant cells, UV signature mutations at a particular site may be serving as a proxy for other causal events resulting from UV exposure, and that determination of clonal expansion phenotypes may underestimate the actual extent because it relies on statistical analysis rather than on studying cells marked with a reporter construct, an approach not possible in humans.

## Materials and Methods

### Experimental design

Human skin samples having a range of sunlight or UV exposures were obtained and their DNA sequenced, in order to quantify early mutational events in skin cancer and assess their usefulness in objectively measuring past UV exposure and predicting future skin cancer risk. High-fidelity DuplexSeq DNA sequencing was used in order to avoid spurious mutations arising from DNA lesions on the original template, thereby allowing low-abundance mutations to be measured. Bioinformatics analyses were conducted to determine the extent of clonal expansion of mutant cells.

### Study subjects and tissue collection

After informed consent, de-identified tissue and saliva were obtained under protocols to D.E.B. approved by the Yale Human Investigation Committee Institutional Review Board and were handled as a biohazard until DNA had been isolated. Anonymized fresh neonatal foreskin and de-identified frozen normal skin from melanoma patients were obtained from the Specimen Resource Core of the Yale SPORE in Skin Cancer (Dr. R. Halaban, co-Director and A. Bacchiocchi, manager), acquired under its IRB protocols. Saliva was collected using the Oragene saliva collection kit (DNA Genotek, Cat OGR-500). Skin discard tissue >1 cm from the margin of a melanoma excision was stored in RNAlater Stabilization Solution (ThermoFisher Scientific, Cat. AM7022) for one week. Other skin samples were obtained as 4 mm punch biopsies or using the STAMP surfactant procedure (*70*). Tissues from patients and volunteers included adult buccal epithelia; skin from the leg or forearm having typical ambient sunlight exposure; and buttock or forearm of patients with cutaneous T-cell lymphoma (CTCL) therapeutically exposed to narrowband UVB (nbUVB) (**Table S1**).

### Separation of epidermis from dermis

Skin biopsies were rinsed in 1X PBS to remove traces of transport medium. The underlying hypodermis was trimmed off as much as possible. For punch biopsies and the melanoma patient (YUDOBI), 2.2-2.5 mL dispase (Zen Bio Cat. CnT-DNP-10, resuspended according to manufacturer’s instructions) was added to a 35 mm dish and the biopsy placed in the solution, with the dermal side submerged. Tissue was incubated at 4°C for 12-16 hr. After incubation, the epidermal layer was carefully teased apart from the underlying layers. The separated epidermis was rinsed in 1X PBS before genomic DNA isolation.

### STAMP (Surfactant-based Tissue Acquisition for Molecular Profiling)

The STAMP procedure was carried out as previously described (*70*), with modifications. For each skin sample, two sterile felt-tip applicator pens (holder #C125 and anti-leak tip #P2, Sunny Packaging, Hefei, Anhui, China), are pre-labeled as “wet” and “dry”. The selected skin area is wiped with 70% alcohol, allowed to dry, and shaved if hair is present. Using a sterile disposable wooden applicator (Dynarex, Thomas Scientific #21A00A189), ∼50 μL of DXB gel [0.25% Laureth-4 (Brij-30 and polyoxyethylene 4 lauryl ether); 0.25% DPS-30 (N-decyl-N,N-dimethyl-3-ammonio-1-propanesulfonate); 15% boric acid/potassium chloride buffer, pH 9.5; 0.25% polysorbate-20, NF grade; 0.5% hydroxypropylcellulose, NF; 0.5% diazolidinyl urea; 0.25% phenylethyl alcohol] Clearista Retexturizing Gel (Skincential Sciences, San Francisco) is applied to the flat surface of the “wet” pen. The skin is stretched and coated with DXB, then firmly rubbed with the forward edge of the DXB-coated pen tip, first in a circular motion for 1 min and then in a linear motion for 1 min. Reaching the dermal-epidermal junction is confirmed by a red speckling pattern; the dermis is not sampled. Residual gel on the skin is recovered by applying the “dry” pen, following the aforementioned steps but with adjustments in time or vigorousness as needed for each patient. The patient sensation is akin to a rug burn and healing occurs in ten days without scarring.

Each wet or dry pen is subsequently placed in a 15 mL Falcon tube containing 500 μL 1X Ca/Mg-free PBS, pH 7.4 (Gibco #10010-023) and frozen at -80°C for 2 hr. (Pens may be stored at -80°C for later processing). After thawing at room temperature for ≤30 min, 30 μL proteinase K and 500 μL buffer AL buffer are added from the DNAeasy Blood & Tissue Kit (Qiagen #69504). Samples are incubated at 56°C for 12-16 hr, after which each lysate is transferred into an eppendorf tube containing 40 μL of 20 mg/mL RNAse A prepared from 100 mg/mL RNAse A (Qiagen, Cat# 19101). Using separate scalpel blades (Integra Miltex Sterile Safety Scalpels, Fisher Scientific, #12460462), each felt tip is shaved and collected into its respective lysate. Following incubation with RNAse A for 5-10 min at room temperature, the tubes are centrifuged at 20,000 g for 13-15 min at room temperature and each supernatant is transferred into a fresh eppendorf tube. The pelleted shavings from the wet and dry felt tips are added to a single QIAshredder column (Qiagen #79654), centrifuged at 20,000 g for 3 min, and the eluate transferred into either of the sample’s supernatants collected earlier. One-half volume of ethanol is added to each of the two supernatants, which are then vortexed and successively passed through a single DNAeasy Blood & Tissue Kit column. Centrifugation and elution are as per manufacturer’s instructions. Purified DNA is quantified using the Qubit 1X dsDNA HS Assay Kit (Invitrogen, #Q33230).

### Primary human keratinocytes

Neonatal human epidermal keratinocytes (NHEK-Neo) pooled from 3 donors (Lonza, Cat. 00192906) were cultured in KBM Gold keratinocyte basal medium (Lonza, Cat. 00192151) supplemented with KGM Gold SingleQuots (Lonza, Cat. 00192152) (hydrocortisone, transferrin, epinephrine, hEGF, insulin, bovine pituitary extract, gentamicin, and amphotericin B) and incubated at 37°C in 5% CO_2_.

#### nbUVB source

The narrowband UVB source (311 nm) consisted of two 20-watt UVB narrowband UVB lamps (Philips TL20W/01), delivered at a fluence rate of 1.29-1.31 J/m^2^/s measured by UVB meter (National Biological Corp. UVB-500C), adjusting the irradiation time to achieve the desired dose.

### Mutations from chronic low-dose UV exposure

To simulate mutations made by chronic low-dose UV or sunlight exposure, keratinocytes starting at passage 5 were exposed to fluences of 0, 300, or 1000 J/m^2^ nbUVB, for a total of 6 to 8 irradiations over a span of 28-33 days (**Fig. S1**). NHEK-Neo cells were seeded in 100 mm culture dishes (Falcon, Cat. 353003) at a cell density of 3,500 cell/cm^2^ and after reaching 30% confluence were irradiated in 2 mL of cold DPBS (Gibco, Cat. 14190144). Cells were allowed to grow after each irradiation, and were collected by trypsinization (Lonza, Cat. CC-5012) at 70-80% confluence, 1-4 days after irradiation depending on the dose-dependent growth rate. Trypsin was neutralized with TNS (Lonza, Cat. CC-5002), cells were centrifuged at 200 x g (Eppendorf 5702), the supernatant was discarded, and the cells were resuspended in HEPES buffered saline (Lonza, Cat. CC-5024). Cells were split into two portions. For passaging, one portion was centrifuged at 200 x g, supernatant was removed, and cell pellet was resuspended in KBM medium and seeded in 100 mm dishes at a cell density of 3,500 cells/cm^2^. For mutation analysis, the remaining cells were centrifuged at 200 x g, washed with PBS, and collected by centrifugation at 4000 x g; the supernatant was discarded and cell pellets were snap frozen in dry ice and stored at -80°C until DNA isolation and mutation detection. Cells irradiated at 0 or 300 J/m^2^ were collected 8 times over 33 days, while cells irradiated at 1000 J/m^2^ were collected at 6 times over 28 days due to lack of growth after the later irradiations.

### DNA isolation

Skin biopsies were cut into small pieces and processed with the Monarch Genomic DNA Purification Kit (NEB, Cat. T3010S) or DNeasy blood and tissue kit (Qiagen, Cat. 69506) according to the manufacturer’s protocol including Proteinase K (Qiagen, Cat. 19131) plus RNase A (Qiagen, Cat. 19101, lot 169047789). DNA from saliva was isolated using prepIT-L2P reagent (DNA Genotek, Cat PT-L2P-1.5) following manufacturer’s instructions. Cell culture pellets were resuspended in 200 μL of DPBS (Gibco, Cat. 14190144) and processed with the DNeasy blood and tissue kit as above, except elution of genomic DNA was in buffer TE (AmericanBio, Cat. AB14033). DNA concentration was measured using the Qubit 1X dsDNA HS Assay Kit (Invitrogen, Cat. Q33230).

### High fidelity Duplex Sequencing to detect low-frequency mutations

Direct DNA sequencing has a Variant Allele Frequency (VAF) detection limit of ∼0.5% per bp due to deamination of cytosines or other DNA damage such as 8-oxo-dG on one strand of the initial duplex, or due to PCR errors in the first PCR cycle. This sensitivity limit can be reduced to 10^-7^ per base or lower by Duplex Sequencing, in which the initial top and bottom strands of the duplex are separately barcoded and their PCR progeny sequenced (*42, 71*). Duplex Sequencing was carried out using the TwinStrand Duplex Sequencing Kit according to manufacturer’s protocol (Rev 1.1, TwinStrand Biosciences, Seattle, WA). Briefly, 0.5-1 μg of genomic DNA was fragmented to ∼300-350 bp ultrasonically (E220 Focused-ultrasonicator, Covaris LLC., Woburn, MA) or to 250-350 bp using fragmentase supplied by TwinStrand, end-repaired and A-tailed, and ligated using the kit-supplied Duplex adapters, then treated with the supplied Library Conditioning Mixture. The libraries were amplified using primers containing dual-index sequences followed by dual hybrid capture with one of several TwinStrand Custom Probe Panels (**Table S4**). These panels consisted of 120 bp probes targeting 43-48 specific genome regions, totaling 13.6-15.4 kb. The probe list was designed using the GRCh38/hg38 reference genome. Indexed libraries were sequenced by the Yale Center for Genomic Analysis using 2×150 bp paired-end sequencing on an Illumina NovaSeq 6000 platform. Run scale was usually 1-5% of a lane, yielding a Duplex Consensus Sequencing coverage depth of typically 15,000-30,000 and thus capable of detecting a VAF of 3-7 ×10^-5^ per nt and MF of ∼4×10^-9^ per nt.

### IndelCorrected-DCS (IC-DCS) alignment pipeline for accurate measurement of mutation recurrence frequencies

Accurate measurement of mutation recurrence frequencies requires not only accurate sequencing but also correct one-to-one alignment with the reference sequence. The IC-DCS pipeline was developed in house to provide correct one-to-one alignment of paired-end sequences derived from indexed DuplexSeq libraries, eliminating the conventional need for local re-alignment (*44, 45*) and optimizing the accuracy of measured mutation recurrence frequencies. Correct one-to-one alignment is achieved by customized use of the standalone BLAT mapping tool. Particularly important is the use of BLAT output psl files to select read pairs that both have unique full-length mapped locations within the same probe region. Most important is the use of axt files, derived from BLAT psl files, to bring all of the selected paired read sequences into correct one-to-one alignment with the DNA reference sequence by removing the positions for any called insertions and by retaining the positions for any called deletions. A “paired-end IC-DCS” is defined here as a pair of single-end IC-DCSs constructed from a family of >2 PCR progeny from each strand of a single parental paired-end duplex, with each family member consisting of a correctly structured read pair sharing the same start, end, and CIGAR string and having unique full-length mapping onto the probe’s contiguous region of interest (including 250 bp flanks). Every overlapping pair of single-end IC-DCSs is replaced by a single consensus IC-DCS. The single-nt and multi-nt variant counts (snv and mnv) are determined without realignment, by summing the variants carried by the covering IC-DCSs. Because oxidative DNA damage is often accompanied by insertion-deletion mutations, this approach also significantly improved calling of putative oxidatively induced mutations. Precise term definitions are provided in **Table S5**.

### Statistical analysis

#### Clonal expansion by allele imbalance

Recurrent mutations resulting from independent stochastic events, such as high CPD frequencies or high C deamination rates, would be equally distributed between a sample’s two alleles; dependent events such as clonal expansion can cause an allele imbalance. Patients carrying a heterozygous single-nucleotide polymorphism (SNP) within 50 nt of a hotspot C→T mutation were examined for statistically significant imbalance as quantified by the binomial test.

#### Clonal expansion by Mahalanobis distance

In cases where heterozygous SNPs were not available (notably the tumor suppressor genes), we used the fact that independent stochastic mutational events at a CC CPD would give the same proportion of 5’C→T:CC→TT:3’C→T in each patient, although that ratio will vary from site to site. Outliers indicate clonal expansion. This three-dimensional comparison was performed using Mahalanobis distances (*72*), applicable when multiple variables are linearly related. Distances from the centroid of the data are statistically independent and have a known chi-square probability distribution, so outliers are observations that break the linearity. Here, different patients will have different total counts due to different UV exposures, with independent events creating a roughly linear centroid in which the proportions remain constant. The analysis was performed using the “robust” option of the program Moutlier in the R package chemometrics (https://CRAN.R-project.org/package=chemometrics), calling the function covMcd in the R package robustbase (https://CRAN.R-project.org/package=robustbase).

The mathematical theory of Mahalanobis distances is based on normally distributed continuous variables, whereas we calculated Mahalanobis distances based on Poisson distributed discrete counts. We believe the robust Moutlier calculations reliably identified outliers, because: a) Unlike measured continuous values, which generally have unknown (and not necessarily approximately normal) distributions, our observed CC mutation counts have Poisson distributions which are necessarily approximately normal for counts >20. b) The robust Moutlier results clearly identified two classes of CC mutation profiles: linearly scaled non-outlier profiles, with robust Mahalanobis distances well below the chi-square threshold for statistical significance; ii) nonlinearly distorted outlier profiles, with robust Mahalanobis distances well above the chi-square threshold. Consequently, classifying CC mutation profiles is highly resistant to inaccuracies in calculated distances and thresholds.

## Acknowledgments

We thank Dr. Ruth Halaban and Ms. Antonella Bacchiocchi for banked skin samples, Dr. Russ Lebovitz for providing large quantities of DBX gel, and Dr. Abhijit Patel for technical advice.

## Author contributions

Conceptualization: DEB

Methodology: VM, MP, KYT, MG, DEB

Patient Recruitment and Biopsy: SN, BC, LF, CK, AB, RH, KYT, MN, SK, MG

Investigation: VM (*in vivo*, DuplexSeq), AGR (*in vitro*)

Bioinformatics: KK Writing:

DEB, VM

## Data and materials availability

All processed data are available in the main text or the supplementary materials. Raw sequencing files (FASTQ) and custom bioinformatics code will be deposited in a publicly available database.

