## Supplemental File for "Pervasive Induction of Regulatory Mutation Microclones in Sun-exposed Skin"

### Supplementary Methods

| Sample ID | Body Site | Age | Gender | Sun Exposure | Collection Method |
|---|---|---|---|---|---|
| FS 2A | foreskin | neonatal | M | none | surgical excision |
| FS 2B | foreskin | neonatal | M | none | surgical excision |
| FS 3A | foreskin | neonatal | M | none | surgical excision |
| FS 3B | foreskin | neonatal | M | none | surgical excision |
| DTC | blood (non-smoker) | 18 | M | transient ambient transdermal | blood draw |
| Saliva 1 | oral epithelium | young adult | M | none | Oragene saliva collection kit |
| Saliva 4 | oral epithelium | young adult | M | none | Oragene saliva collection kit |
| Saliva 2 | oral epithelium | older adult | F | none | Oragene saliva collection kit |
| Saliva 3 | oral epithelium | older adult | M | none | Oragene saliva collection kit |
| Y008 | Right Buttock (CTCL) | 55 | M | nbUVB | 4mm punch |
| Y0011 | Right Buttock (CTCL) | 59 | M | nbUVB | 4mm punch |
| Y0015 | Outer, Lower Left Arm (AM) | 62 | F | ambient | 4mm punch |
| Y0016 | Outer, Lower Left Arm (AM) | 59 | M | ambient | 4mm punch |
| Y0017 | Outer Left Arm, Between Wrist and Elbow (AM) | 70 | F | ambient | 4mm punch |
| SY107 | Back, Right Upper Shoulder (CTCL) | 59 | M | nbUVB | STAMP |
| SY102 | Right Arm, Outside Arm, Lower (CTCL) | 56 | M | nbUVB | STAMP |
| SY114 | Right Distal Forearm (CTCL) | 68 | M | nbUVB | STAMP |
| SY118 | Outer Right Forearm (CTCL) | 55 | M | nbUVB | STAMP |
| SY112 | Outer Right Arm Just Under Elbow (TPN) | 72 | M | - | STAMP |
| SY124 | Outer Right Arm Just Above Elbow (AM) | 70 | F | ambient | STAMP |
| SY126 | Outer Right Arm Just Above Elbow (AM) | 56 | F | ambient | STAMP |
| SY132 | Outer Right Arm Between The Elbow And Wrist (AM) | 49 | F | ambient | STAMP |
| SY130 | Outer Right Arm Between Wrist And Elbow (AM) | 59 | F | ambient | STAMP |
| SY138 | Left Arm Closer To Elbow (AM) | 67 | F | ambient | STAMP |
| SY146 | Outer Left Arm Between Wrist And Elbow (AM) | 55 | M | ambient | STAMP |
| DNA-63 | Leg (phototype 3, asian)<br>Non Sun-Exposed; hardly any sun-exposure | 50 | M | intermittent ambient | STAMP |
| UM001 | Left Posterior Mid Forearm (AM) | 76 | M | ambient | 4mm punch |
| UM002 | Right Distal Forearm (AM) | 49 | F | ambient | 4mm punch |
| UM003 | Right Distal Forearm (AM) | 46 | F | ambient | 4mm punch |
| UM005 | Right Proximal Forearm (AM) | 71 | F | ambient | 4mm punch |
| YUDOBI | ipsilateral neck (tumor left cheek) | 78 | M | extensive sun exposure in youth (caddy, suntanning), with non-blistering sunburns; prior SCCs, BCCs, melanoma in situ; several melanomas; family history of various CAs | surgical excision |

### Table S1. Patient characteristics.

MutationFrequenciesSummary

| | Independent mutations | MFminI no exposure | MFminI low exposure | MFminI high exposure | MFmaxI no exposure | MFmaxI high exposure | VAFminI low exposure | VAFminI high exposure | VAFmaxI low exposure (incl recurrent so ~MFmaxI) | VAFmaxI high exposure (incl recurrent so ~MFmaxI) | Excess Recurrence (wrt GenoTox avgVAF) | CPD-unexplainable Excess Recurrence (prev column left if CPD HH ~100x–500x genome avg) | Mutations per Cell (maxI) | Source | Deductions |
|---|---|---|---|---|---|---|---|---|---|---|---|---|---|---|---|
| **Literature** (any base change) | | | | | | | | | | | | | | | |
| Cell culture colonies, UV *above spontaneous* , reporter, any bc estimate assumes 100 target nt in gene | $10^{-7}$–$10^{-5}$ (low –> hi dose) | – | $10^{-7}$ | $10^{-5}$ | – | $10^{-5}$ | $10^{-7}$ | $10^{-5}$ | | $10^{-5}$ | – | – | 600 – 60,000 | Menon '23 | |
| Sun-exposed skin, CA genes, any base change | – | – | – | – | – | $0.4x10^{-5}$ | – | – | – | – | – | – | 24,000 | Martincorena '15 | Small samples give hi VAF, so avoids spurious clones to the extent that >1% VAF are non-spurious (cf Valentine et al. 2020); counted any mut not different-muts, so is MFmaxI |
| | | | | | | | | | | | | | | | So in the 2015 paper, repeated exposures +clonal expansion gives a *recurrent* MFmaxI (0.4E-05) that is less than our 1.0E-05 estimate of *independent* pre-clonal founders (MFminI) arising from a single hi UV dose to a reporter gene in cell culture. Thus either: a) the skin's independent founder MFminI is actually at the lower end of our single exposure estimate (10^-7 ) OR b) their focus on clones above 1% VAF underdetects mutations. Or both. |
| **DuplexSeq** (**TwinStrand**, present custom capture panels) | | | | | | | | | | | | | | | |
| Present detection limit (fcn of # genomes and, for MF, panel kb) | | *$4x10^{-9}$* | *$4x10^{-9}$* | *$4x10^{-9}$* | *$4x10^{-9}$* | *$4x10^{-9}$* | *$4x10^{-5}$* | *$4x10^{-5}$* | – | *$4x10^{-5}$* | | | | | |
| ***Universal targets*** (all probe nt) | | | | | | | | | | | | | | | |
| Neonatal foreskin, univ targets, any base change | | $0.7x10^{-5}$ | | | $1.5x10^{-5}$ | | | | | | | | 90,000 | Fig 4 | |
| Sun-exposed skin, univ targets, any base change | – | – | | $1x10^{-5}$ | – | $4x10^{-5}$ | – | – | – | – | – | – | 240,000 | Fig 4 | So DuplexSeq any-base-change MFmaxI is 10x Martincorena 2015. So 10x of line 7 discrepancy comes from (b) underdetection. |
| Neonatal foreskin, univ targets, UV signature | – | *~$4x10^{-9}$* | | | *~$4x10^{-9}$* | | | | | | | | 24 | Fig 4 | So 10,000x differential between sun-shielded and sun-exposed. |
| Sun-exposed skin, univ targets, UV signature | – | – | | **$0.2x10^{-5}$** | **–** | $3x10^{-5}$ | – | – | – | – | – | – | 180,000 | Fig 4 | So DuplexSeq UV signature MFminI is in mid-range of estimated. Thus 10x of line 7 discrepancy comes from (a) modest dose.<br>Still, our DuplexSeq value is also less than one would expect from multiple exposures. MFmaxI is 10x this on average. |
| ***Univ targets split, sun-exposed skin, UV signature*** **regions –** | | | | | | | | | | | | | | | |
| GenoTox region (UV signature) | – | – | | **$0.2x10^{-5}$** | **–** | $0.5x10^{-5}$ | – | – | – | – | – | – | – | | So GenoTox MFminI = universal probe MFminI; GenoTox neutral drift (MFmaxI/MFminI) is ~2.5x |
| CA Genes (tumor suppressor + oncogene + mut in sun-exp skin) | – | – | | $0.3x10^{-5}$ | – | $3x10^{-5}$ | – | – | – | – | – | – | – | | So CA gene region muts recur (clonally expand) 10x on average |
| CPD HH regions | | – | | *$05x10^{-5}$ | – | $10^{-4}$ | – | – | – | – | – | – | – | | So CPD HH *region* muts recur 20x on average |
| **nt –** | | | | | | | | | | | | | | | |
| GenoTox region (avgVAF, UV signature) | – | – | – | – | – | – | – | – | – | $10^{-6}$ | 1 | – | 0.01 | Fig 3 | detection limit ~1/845*35,000 = 3E-08 (data fr *summedVAFpct_GenoTox C>T diPyv2.xlsx) |
| TP53 & NOTCH1 tumor hotspot nt | – | – | < $2x10^{-7}$ | $3x10^{-5}$ | – | – | – | – | – | 0 or $10^{-3}$–$10^{-1}$ | 0 or 100 – 10,000<br>200 – 20,000 using the more sensitive MFmaxI | – | – | Fig 3, some biopsies; summed VAFpctC>T diPy ROI bugfix reorder universal probes | So TS genes are sporadic genomic dosimeters. Some patient TS VAFs are 200-20,000x GT MFmaxI (ie ratio to GT founders expanded by neutral drift). Most biopsies are 0. |
| CPD HH nt | – | – | $5x10^{-6}$ | > $4x10^{-5}$ | – | – | – | – | – | **$10^{-3}$–$10^{-1}$** | 100 – 10,000<br>or 200 – 20,000 | 1x – 40x | 17* | Fig 3, ~all biopsies; summed VAFpctC>T diPy ROI bugfix reorder universal probes | So CPD HH are ubiquitous genomic dosimeters. All patient CPD HH VAFs are 200-20,000x GT MFmaxI. **Upper range is more than CPD excess.** |
| | | | | | | | | | | | | | * Using the midrange value of 10^-2 and ~2000 CPD hyperhotspots in keratinocytes (Garcia-Ruiz et al. 2022) | | |
| | | | | | | | | | | | | | GenoTox probe region length: 2.4 kb | | |
| ***Chronic expt*** (C>TdiPy, so UV signature +deamination) | | | | | | | | | | | | | | | |
| GenoTox region (avgVAF) | – | – | – | – | – | – | | | low 1st $1.4x10^{-7}$<br>low 4th 1.1x10^-6<br>low 6th 1.4x10^-6 | **hi 1st 5.6x10^-7**<br>hi 4th 2x10^-6<br>hi 6th 9.5x10^-7 | 1 | – | 1st: GenoTox 0.0003–0.001 genome 850 – 3500 6th: GenoTox 0.003 genome 7000 | Fig 6 (low = 300 J/m2 nbUVB high = 1000) | So the low-dose single irradiation GenoTox UV VAF (1.4E-07) agrees with our estimate from literature; hi dose is in mid-range.<br>This single-irradiation GT UV VAF is ~1/200 the UV MFmaxI we see in skin (3E-05). Is skin due to repeat exposures or clones? |
| GenoTox hot nt | – | – | – | – | – | – | | | low 1st $5x10^{-5}$ [1 of 4]<br>low 4th 5x10^-4 [1 of 4]<br>low 6th 3x10^-4 [3 of 4] >6x incl all 4 | **hi 1st 5x10^-5** [3 of 4]<br>hi 4th 1x10^-4 [2 of 4]<br>hi 6th 10^-3 [1 of 4] <20x for all 4 | – | – | – | Fig 6 | So at chronic expt's 1st hi dose, 3 of 4 GT hot sites (5E-05) were 350x the MF of GT ordinary sites (1.4E-07) and just above our TwinStrand VAF detection limit. |
| Cancer gene, skin clone gene nt | – | – | – | – | – | – | | | | 0 | unmeasurable | – | – | Fig 6 | So oncogenes and some tumor suppressor genes are below our VAF detection limit; so we don't know their actual VAFminI wrt our estimated VAFminI. |
| TP53, NOTCH1 tumor hotspot nt | – | – | – | – | – | – | | | low 1st, 6th 0 | hi 1st 5x10^-5 [2 of 48] then disappear<br>hi 4th ~6x10^-5 [5 of 48] then disappear<br>hi 6th 0 | unmeasurable | – | – | Fig 6 | So at low dose, TP53 and NOTCH1 tumor hotspot nt are like the GT non-hot sites, below detection limit.<br>At high dose, 5 of 48 TS tumor hotspot nt are like the GT hot sites; remainder are like ordinary GT and oncogenes.<br>So no evidence that TS gene mutations rapidly and autonomously expand over 30 days. So lack of evidence of expansion in CPD HH would not be conclusive. |
| CPD HH nt | – | – | – | – | – | – | | | low 1st 6x10^-5 – 6x10^-4 [19 of 30 | – | 429 – 4290 | 1x – 8x | – | Fig 6 | At the low end, initial CPD HH VAFs are 400x the GT UV avgVAF (1.4E-07), and higher than TP53, and NOTCH1. At the hi end, they are 4000x. |
| CPD HH nt | – | – | – | – | – | – | | | low 4th 3x10^-4** – 3x10^-3 [25 of 30 | – | 273 – 2730 | 1x – 5x | – | Fig 6 **for those present at 1st | 4x low exposure -> ~5x UV VAFmaxI, so linear w exposures (time & dose) |
| CPD HH nt | – | – | – | – | – | – | | | low 6th 3x10^-4** – 10^-2 [25 of 30 | – | 214 – 7143 | 1x – 15x | – | Fig 6 **for those present at 1st | 6x low exposure -> ~5x – 15x UV VAFmaxI of 1st; highest are lines on log plot, so exponential, logy = a+kt y=10^(a+kt) = a'*10^kt; non-log graph shows highest are exponential.<br>So 6 low exposures or 30 days allow exponential growth of some CPD HH. |
| CPD HH nt | – | – | – | – | – | – | | | – | hi 1st 6x10^-5 – **10^-3** [24 of 30] | 107 – 1786 | 1x – 4x | 0.4* | Fig 6 | Initial 100x – 2000x GT hi UV avgVAF. High dose 3.3x low dose gives 1-2x mutations, so sublinear w dose at 1st irrad. |
| **CPD HH nt** | – | – | – | – | – | – | | | – | hi 4th 3x10^-4** – 3x10^-3 [30 of 30] | 150 – 1500 | 1x – 3x | 2* | Fig 6 | 4x hi exposure -> 3-5x VFmaxI at 1st irradiation, so linear w exposures; 5 – 50x tumor suppressor genes at 4th irradiation |
| CPD HH nt | – | – | – | – | – | – | | | – | hi 6th 6x10^-5** – 5x10^-2 [25 of 30] | 65 – 52,632 | 1x – 100x | 4* | Fig 6 **for those present at 1st | 6x hi exposure -> 1x – 50x mutations at 1st irradiation; highest are lines on log plot, so exponential; non-log graph shows highest are exponential. So 6 hi exposures or 30 days allow exponential growth of some CPD HH. |

any bc = any base change type

Note: MFminI will be an underestimate if same site is hit recurrently, esp CPD HH.

**universal** = probe panel used in all expts

**UV signature** = C->TdiPy w deamination subtracted

HH = hyperhotspot

### Table S2

**Table S3 Allelic imbalance of SNP-adjacent mutations**

| Patient | chr | location (mutation & SNP if alt allele) | SNP location | Mutation | Gene | Site Property | | SNP allele | Ref allele | Min | Max | Ratio | Sum | Min | Sum | P | -logP |
|---|---|---|---|---|---|---|---|---|---|---|---|---|---|---|---|---|---|
| **Tumor suppressor gene** | | | | | | | | | | | | | | | | | |
| SY112 | chr17 | 7673523#7673533 | 7673523 | AA>GT_1 | TP53 | | 9 0 | 9 | 0 | 0 | 9 | INF | 9 | | 9 | 3.91E-03 | 2.4 |
| SY112 | chr17 | 7673534#7673535 | 7673523 | CC>TT_1 | TP53 | | 0 9 | 0 | 9 | 0 | 9 | INF | 9 | | 9 | 3.91E-03 | 2.4 |
| SY112 | chr17 | 7674775 | 7674797 | G>A_1 | TP53 | | 0 5 | 0 | 5 | 0 | 5 | INF | 5 | | 5 | 6.25E-02 | 1.2 |
| SY112 | chr17 | 7675322#7675358 | 7675322 | AT>GG_2 | TP53 | | 5 8 | 5 | 8 | 5 | 8 | 1.6 | 13 | | 13 | 5.81E-01 | 0.2 |
| SY132 | chr17 | 7674858 | 7674892 | C>T_2 | TP53 | | 10 2 | 10 | 2 | 2 | 10 | 5.0 | 12 | 2 | 12 | 3.86E-02 | 1.4 |
| SY132 | chr17 | 7674875#7674892 | 7674892 | GT>AC_2 | TP53 | | 4 0 | 4 | 0 | 0 | 4 | INF | 4 | 0 | 4 | 1.25E-01 | 0.9 |
| SY132 | chr17 | 7674878 | 7674892 | A>C_2 | TP53 | | 0 5 | 0 | 5 | 0 | 5 | INF | 5 | 0 | 5 | 6.25E-02 | 1.2 |
| SY132 | chr17 | 7674891#7674892 | 7674892 | GT>AC_2 | TP53 | | 21 0 | 21 | 0 | 0 | 21 | INF | 21 | 0 | 21 | 9.54E-07 | 6.0 |
| SY132 | chr17 | 7674892#7674899 | 7674892 | TG>CA_2 | TP53 | | 11 0 | 11 | 0 | 0 | 11 | INF | 11 | 0 | 11 | 9.77E-04 | 3.0 |
| SY114 | chr17 | 7674858 | 7674892 | C>T_1 | TP53 | | 0 24 | 0 | 24 | 0 | 24 | INF | 24 | | 24 | 1.20E-07 | 6.9 |
| SY114 | chr17 | 7674924 | 7674892 | C>A_1 | TP53 | | 0 7 | 0 | 7 | 0 | 7 | INF | 7 | | 7 | 1.60E-02 | 1.8 |
| SY107 | chr17 | 7674044#7674089 | 7674089 | TA>GC_1 | TP53 flank | | 17 18 | 17 | 18 | 17 | 18 | 1.1 | 35 | 17 | 35 | 1.00E+00 | 0.0 |
| SY112 | chr17 | 7685993#7686030 | 7685993 | GA>CC_1 | TP53 flank | | 2 4 | 2 | 4 | 2 | 4 | 2.0 | 6 | | 6 | 6.88E-01 | 0.2 |
| **CPD hyperhotspots** | | | | | | | | | | | | | | | | | |
| SY146 | chr6 | 80004496#80004519 | | GC>CT_2 | TTK | HH | 5 2 | 5 | 2 | 2 | 5 | 2.5 | 7 | | 7 | 4.50E-01 | 0.3 |
| SY146 | chr6 | 80004519 | | C>T_3 | TTK | HH | 0 7 | 0 | 7 | 0 | 7 | INF | 7 | | 7 | 1.60E-02 | 1.8 |
| Y0017 | chr6 | 80004496#80004519 | 80004496 | GC>CT_1 | TTK | HH | 7 5 | 7 | 5 | 5 | 7 | 1.4 | 12 | | 12 | 7.74E-01 | 0.1 |
| Y0017 | chr6 | 80004496#80004530 | 80004496 | GG>CA_1 | TTK | | 5 0 | 5 | 0 | 0 | 5 | INF | 5 | | 5 | 6.25E-02 | 1.2 |
| Y0017 | chr6 | 80004516#80004530 | 80004516 | GG>TA_2 | TTK | | 3 2 | 3 | 2 | 2 | 3 | 1.5 | 5 | | 5 | 1.00E+00 | 0.0 |
| Y0017 | chr6 | 80004519 | 80004516 | C>T_2 | TTK | HH | 1 8 | 1 | 8 | 1 | 8 | 8.0 | 9 | | 9 | 3.91E-02 | 1.4 |
| SY102 | chr6 | 80004496#80004519 | 80004496 | GC>CT_1 | TTK | HH | 10 10 | 10 | 10 | 10 | 10 | 1.0 | 20 | 10 | 20 | 1.18E+00 | -0.1 |
| SY102 | chr6 | 80004519 | 80004516 | C>T_2 | TTK | HH | 1 15 | 1 | 15 | 1 | 15 | 15.0 | 16 | 1 | 16 | 5.19E-04 | 3.3 |
| SY107 | chr6 | 80004496#80004519 | 80004496 | GC>CT_1 | TTK | HH | 102 1 | 102 | 1 | 1 | 102 | 102.0 | 103 | 1 | 103 | 2.05E-29 | 28.7 |
| SY107 | chr6 | 80004496#80004523 | 80004496 | GC>CT_1 | TTK | HHW | 5 3 | 5 | 3 | 3 | 5 | 1.7 | 8 | 3 | 8 | 7.27E-01 | 0.1 |
| SY107 | chr6 | 80004516#80004519 | 80004516 | GC>TT_2 | TTK | HH | 28 79 | 28 | 79 | 28 | 79 | 2.8 | 107 | 28 | 107 | 8.42E-07 | 6.1 |
| SY118 | chr6 | 80004496#80004519 | 80004496 | GC>CT_1 | TTK | HH | 16 33 | 16 | 33 | 16 | 33 | 2.1 | 49 | 16 | 49 | 2.13E-02 | 1.7 |
| SY118 | chr6 | 80004516#80004519 | 80004516 | GC>TT_2 | TTK | HH | 3 50 | 3 | 50 | 3 | 50 | 16.7 | 53 | 3 | 53 | 5.52E-12 | 11.3 |
| SY118 | chr6 | 80004515 | 80004516 | G>A_2 | TTK | | 0 9 | 0 | 9 | 0 | 9 | INF | 9 | 0 | 9 | 3.91E-03 | 2.4 |
| SY114 | chr6 | 80004475#80004519 | 80004475 | CC>GT_1 | TTK | HH | 19 16 | 19 | 16 | 16 | 19 | 1.2 | 35 | | 35 | 7.40E-01 | 0.1 |
| SY114 | chr6 | 80004496#80004519 | 80004496 | GC>CT_2 | TTK | HH | 4 29 | 4 | 29 | 4 | 29 | 7.3 | 33 | | 33 | 1.10E-05 | 5.0 |
| SY114 | chr6 | 80004519#80004541 | 80004541 | CA>TG_3 | TTK | HH | 8 25 | 8 | 25 | 8 | 25 | 3.1 | 33 | | 33 | 4.60E-03 | 2.3 |
| SY114 | chr6 | 80004519#80004559 | 80004559 | CC>TT_4 | TTK | HH | 2 18 | 2 | 18 | 2 | 18 | 9.0 | 20 | | 20 | 4.00E-04 | 3.4 |
| SY132 | chr5 | 138543244#138543258 | 138543258 | CG>TA_1 | ETF1 | | 4 0 | 4 | 0 | 0 | 4 | INF | 4 | 0 | 4 | 1.25E-01 | 0.9 |
| SY132 | chr5 | 138543258#138543281 | 138543258 | GG>AA_1 | ETF1 | HH | 15 9 | 15 | 9 | 9 | 15 | 1.7 | 24 | 9 | 24 | 3.07E-01 | 0.5 |
| SY132 | chr5 | 138543258#138543281#138543282 | 138543258 | GGG>AAA_1 | ETF1 | HH, HH | 7 2 | 7 | 2 | 2 | 7 | 3.5 | 9 | 2 | 9 | 1.80E-01 | 0.7 |
| SY132 | chr5 | 138543258#138543282 | 138543258 | GG>AA_1 | ETF1 | HH | 35 124 | 35 | 124 | 35 | 124 | 3.5 | 159 | 35 | 159 | 7.16E-13 | 12.1 |
| SY132 | chr5 | 69189491 | 69189539 | G>A_1 | CENPH | | 1 8 | 1 | 8 | 1 | 8 | 8.0 | 9 | 1 | 9 | 3.91E-02 | 1.4 |
| SY132 | chr5 | 69189499#69189539 | 69189539 | AC>CG_1 | CENPH | | 6 3 | 6 | 3 | 3 | 6 | 2.0 | 9 | 3 | 9 | 5.08E-01 | 0.3 |
| SY132 | chr5 | 69189508#69189539 | 69189539 | GC>AG_1 | CENPH | | 3 2 | 3 | 2 | 2 | 3 | 1.5 | 5 | 2 | 5 | 1.00E+00 | 0.0 |
| SY132 | chr5 | 69189519 | 69189539 | T>G_1 | CENPH | | 29 18 | 29 | 18 | 18 | 29 | 1.6 | 47 | 18 | 47 | 1.44E-01 | 0.8 |
| SY132 | chr5 | 69189528#69189529 | 69189539 | GG>AA_1 | CENPH | HH, HH | 13 4 | 13 | 4 | 4 | 13 | 3.3 | 17 | 4 | 17 | 4.90E-02 | 1.3 |
| SY132 | chr5 | 69189528#69189539 | 69189539 | GC>AG_1 | CENPH | HH | 18 15 | 18 | 15 | 15 | 18 | 1.2 | 33 | 15 | 33 | 7.28E-01 | 0.1 |
| SY132 | chr5 | 69189529 | 69189539 | G>A_1 | CENPH | HH | 19 18 | 19 | 18 | 18 | 19 | 1.1 | 37 | 18 | 37 | 1.00E+00 | 0.0 |
| SY132 | chr5 | 69189539#69189542 | 69189539 | CG>GA_1 | CENPH | | 4 0 | 4 | 0 | 0 | 4 | INF | 4 | 0 | 4 | 1.25E-01 | 0.9 |
| SY132 | chr5 | 69189539#69189559 | 69189539 | CC>GT_1 | CENPH | | 4 0 | 4 | 0 | 0 | 4 | INF | 4 | 0 | 4 | 1.25E-01 | 0.9 |
| Y0016 | chr5 | 69189519 | 69189539 | T>G_1 | CENPH | | 16 7 | 16 | 7 | 7 | 16 | 2.3 | 23 | 7 | 23 | 9.31E-02 | 1.0 |
| Y0016 | chr5 | 69189529 | 69189539 | G>A_1 | CENPH | HH | 1 15 | 1 | 15 | 1 | 15 | 15.0 | 16 | 1 | 16 | 5.19E-04 | 3.3 |
| SY114 | chr5 | 69189508#69189539 | 69189539 | GC>AG_1 | CENPH | | 11 5 | 11 | 5 | 5 | 11 | 2.2 | 16 | | 16 | 2.10E-01 | 0.7 |
| SY114 | chr5 | 69189519#69189539 | 69189539 | TC>GG_1 | CENPH | | 5 3 | 5 | 3 | 3 | 5 | 1.7 | 8 | | 8 | 7.30E-01 | 0.1 |
| SY114 | chr5 | 69189525 | 69189539 | G>A_1 | CENPH | HHW | 4 2 | 4 | 2 | 2 | 4 | 2.0 | 6 | | 6 | 6.90E-01 | 0.2 |
| SY114 | chr5 | 69189528#69189529#69189539 | 69189539 | GGC>AAG_1 | CENPH | HH, HH | 6 7 | 6 | 7 | 6 | 7 | 1.2 | 13 | | 13 | 1.00E+00 | 0.0 |
| SY114 | chr5 | 69189528#69189539 | 69189539 | GC>AG_1 | CENPH | HH | 9 25 | 9 | 25 | 9 | 25 | 2.8 | 34 | | 34 | 3.10E-01 | 0.5 |
| SY114 | chr5 | 69189529#69189539 | 69189539 | GC>AG_1 | CENPH | HH | 16 7 | 16 | 7 | 7 | 16 | 2.3 | 23 | | 23 | 9.30E-02 | 1.0 |
| SY114 | chr5 | 69189539#69189546 | 69189539 | CC>GT_1 | CENPH | | 14 0 | 14 | 0 | 0 | 14 | INF | 14 | | 14 | 1.20E-04 | 3.9 |
| SY112 | chr19 | 5791156#5791181#5791182 | 5791156 | TCC>CTT_1 | DUS3L | HH, HH | 14 1 | 14 | 1 | 1 | 14 | 14.0 | 15 | | 15 | 9.77E-04 | 3.0 |
| SY112 | chr19 | 5791182 | 5791156 | C>T_1 | DUS3L | HH | 2 5 | 2 | 5 | 2 | 5 | 2.5 | 7 | | 7 | 4.53E-01 | 0.3 |
| SY118 | chr19 | 5791156#5791165 | 5791156 | TG>CA_1 | DUS3L | | 6 2 | 6 | 2 | 2 | 6 | 3.0 | 8 | 2 | 8 | 2.89E-01 | 0.5 |
| SY118 | chr19 | 5791181#5791182 | 5791156 | CC>TT_1 | DUS3L | HH, HH | 1 5 | 1 | 5 | 1 | 5 | 5.0 | 6 | 1 | 6 | 2.19E-01 | 0.7 |
| SY112 | chr6 | 151452218#151452246 | 151452218 | GC>CT_2 | RMND1//C6orf211 | HH | 1 7 | 1 | 7 | 1 | 7 | 7.0 | 8 | | 8 | 7.03E-02 | 1.2 |
| SY132 | chr3 | 157174984#157175015 | 157175015 | CT>TC_1 | RNA5SP146//RP11-550I24.2 | | 9 0 | 9 | 0 | 0 | 9 | INF | 9 | 0 | 9 | 3.91E-03 | 2.4 |
| SY132 | chr3 | 157174992#157175015 | 157175015 | GT>AC_1 | RNA5SP146//RP11-550I24.2 | HH | 2 4 | 2 | 4 | 2 | 4 | 2.0 | 6 | 2 | 6 | 6.88E-01 | 0.2 |
| SY132 | chr3 | 157175009#157175015 | 157175015 | GT>AC_1 | RNA5SP146//RP11-550I24.2 | | 5 2 | 5 | 2 | 2 | 5 | 2.5 | 7 | 2 | 7 | 4.53E-01 | 0.3 |
| SY132 | chr3 | 157175012#157175015 | 157175015 | AT>TC_1 | RNA5SP146//RP11-550I24.2 | | 12 1 | 12 | 1 | 1 | 12 | 12.0 | 13 | 1 | 13 | 3.42E-03 | 2.5 |
| SY107 | chr3 | 157174992#157175015 | 157175015 | GT>AC_1 | RNA5SP146//RP11-550I24.2 | HH | 4 1 | 4 | 1 | 1 | 4 | 4.0 | 5 | 1 | 5 | 3.75E-01 | 0.4 |
| SY107 | chr3 | 157175009#157175015 | 157175015 | GT>AC_1 | RNA5SP146//RP11-550I24.2 | | 12 3 | 12 | 3 | 3 | 12 | 4.0 | 15 | 3 | 15 | 3.52E-02 | 1.5 |
| SY146 | chr10 | 89701491#89701511 | 89701511 | TT>GG_1 | RP11-80H5.9//KIF20B | | 5 2 | 5 | 2 | 2 | 5 | 2.5 | 7 | | 7 | 1.9 | -0.3 |
| SY118 | chr10 | 89701495#89701496#89701511 | 89701511 | GGT>AAG_1 | RP11-80H5.9//KIF20B | | 9 0 | 9 | 0 | 0 | 9 | INF | 9 | 0 | 9 | 3.91E-03 | 2.4 |
| SY114 | chr10 | 89701491#89701511 | 89701511 | TT>GG_1 | RP11-80H5.9//KIF20B | | 3 1 | 3 | 1 | 1 | 3 | 3.0 | 4 | | 4 | 6.30E-01 | 0.2 |
| SY114 | chr10 | 89701504#89701511 | 89701511 | CT>TG_1 | RP11-80H5.9//KIF20B | HH | 6 2 | 6 | 2 | 2 | 6 | 3.0 | 8 | | 8 | 2.90E-01 | 0.5 |
| SY114 | chr10 | 89701507#89701511 | 89701511 | CT>TG_1 | RP11-80H5.9//KIF20B | HHW | 6 0 | 6 | 0 | 0 | 6 | INF | 6 | | 6 | 3.10E-02 | 1.5 |
| SY102 | chr3 | 51995944 | 51995975 | G>A_1 | RPL29 | HH | 2 4 | 2 | 4 | 2 | 4 | 2.0 | 6 | 2 | 6 | 6.88E-01 | 0.2 |
| SY102 | chr3 | 51995975#51996017 | 51995975 | GG>AA_1 | RPL29 | HH | 3 0 | 3 | 0 | 0 | 3 | INF | 3 | 0 | 3 | 2.50E-01 | 0.6 |
| SY118 | chr3 | 51995944#51995975 | 51995975 | GG>AA_1 | RPL29 | HH | 17 31 | 17 | 31 | 17 | 31 | 1.8 | 48 | 17 | 48 | 5.95E-02 | 1.2 |
| SY118 | chr3 | 51995947#51995975 | 51995975 | CG>TA_1 | RPL29 | | 9 1 | 9 | 1 | 1 | 9 | 9.0 | 10 | 1 | 10 | 2.15E-02 | 1.7 |
| SY112 | chr1 | 155262283#155262293 | 155262283 | CG>TA_1 | SCAMP3 | HH | 7 0 | 7 | 0 | 0 | 7 | INF | 7 | | 7 | 1.56E-02 | 1.8 |
| SY132 | chr1 | 155262283#155262293 | 155262283 | CG>TA_1 | SCAMP3 | HH | 4 2 | 4 | 2 | 2 | 4 | 2.0 | 6 | 2 | 6 | 6.88E-01 | 0.2 |
| SY132 | chr1 | 155262283#155262293#155262294 | 155262283 | CGG>TAA_1 | SCAMP3 | HH, HH | 3 7 | 3 | 7 | 3 | 7 | 2.3 | 10 | 3 | 10 | 3.44E-01 | 0.5 |
| SY107 | chr1 | 155262283#155262293 | 155262283 | CG>TA_1 | SCAMP3 | HH | 11 2 | 11 | 2 | 2 | 11 | 5.5 | 13 | 2 | 13 | 2.25E-02 | 1.6 |
| SY107 | chr1 | 155262283#155262293#155262294 | 155262283 | CGG>TAA_1 | SCAMP3 | HH, HH | 6 0 | 6 | 0 | 0 | 6 | INF | 6 | 0 | 6 | 3.13E-02 | 1.5 |
| SY107 | chr1 | 155262294 | 155262283 | G>A_1 | SCAMP3 | HH | 1 8 | 1 | 8 | 1 | 8 | 8.0 | 9 | 1 | 9 | 3.91E-02 | 1.4 |
| SY112 | chr3 | 41962206#41962237 | 41962206 | TG>AA_1 | ULK4 | HHW | 24 2 | 24 | 2 | 2 | 24 | 12.0 | 26 | | 26 | 1.05E-05 | 5.0 |
| SY112 | chr3 | 41962206#41962240 | 41962206 | TG>AA_1 | ULK4 | HH | 10 6 | 10 | 6 | 6 | 10 | 1.7 | 16 | | 16 | 4.54E-01 | 0.3 |
| SY114 | chr3 | 41962206#41962214 | 41962206 | TC>AT_1 | ULK4 | | 14 0 | 14 | 0 | 0 | 14 | INF | 14 | | 14 | 1.20E-04 | 3.9 |
| SY114 | chr3 | 41962206#41962246 | 41962206 | TC>AT_1 | ULK4 | | 4 1 | 4 | 1 | 1 | 4 | 4.0 | 5 | | 5 | 3.80E-01 | 0.4 |
| SY114 | chr3 | 41962237 | 41962206 | G>A_1 | ULK4 | HHW | 0 6 | 0 | 6 | 0 | 6 | INF | 6 | | 6 | 3.10E-02 | 1.5 |
| SY112 | chr3 | 124730362#124730387 | 124730362 | AG>GA_1 | UMPS | HH | 15 14 | 15 | 14 | 14 | 15 | 1.1 | 29 | | 29 | 1.00E+00 | 0.0 |
| Y0017 | chr3 | 124730362#124730387#124730444 | 124730362&124730444 | AGA>GAG_1 | UMPS | HH | 5 0 | 5 | 0 | 0 | 5 | INF | 5 | | 5 | 6.25E-02 | 1.2 |
| SY102 | chr3 | 124730362#124730387#124730444 | 124730362&124730444 | AGA>GAG_1 | UMPS | HH | 0 4 | 0 | 4 | 0 | 4 | INF | 4 | 0 | 4 | 1.25E-01 | 0.9 |
| SY107 | chr3 | 124730362#124730387 | 124730362 | AG>GA_1 | UMPS | HH | 15 6 | 15 | 6 | 6 | 15 | 2.5 | 21 | 6 | 21 | 7.84E-02 | 1.1 |
| SY107 | chr3 | 124730362#124730394 | 124730362 | AG>GA_1 | UMPS | | 9 1 | 9 | 1 | 1 | 9 | 9.0 | 10 | 1 | 10 | 2.15E-02 | 1.7 |
| SY107 | chr3 | 124730394#124730444 | 124730444 | GA>AG_2 | UMPS | | 9 0 | 9 | 0 | 0 | 9 | INF | 9 | 0 | 9 | 3.91E-03 | 2.4 |
| SY107 | chr3 | 124730399 | 124730362 | C>T_1 | UMPS | | 1 4 | 1 | 4 | 1 | 4 | 4.0 | 5 | 1 | 5 | 3.75E-01 | 0.4 |
| SY107 | chr3 | 124730399 | 124730444 | C>T_2 | UMPS | | 3 4 | 3 | 4 | 3 | 4 | 1.3 | 7 | 3 | 7 | 1.00E+00 | 0.0 |
| SY114 | chr3 | 124730362#124730387#124730444 | 124730362&124730444 | AGA>GAG_1 | UMPS | HH | 0 11 | 0 | 11 | 0 | 11 | INF | 11 | | 11 | 9.80E-04 | 3.0 |
| Y0017 | chr12 | 14803509 | 14803542 | T>G_1 | WBP11//C12orf60 | | 1 4 | 1 | 4 | 1 | 4 | 4.0 | 5 | | 5 | 3.75E-01 | 0.4 |
| **GenoTox non-selectable region** | | | | | | | | | | | | | | | | | |
| Y0017 | chr18 | 5751557#5751576 | 5751576 | GT>AC_2 | Genotox-TS | | 6 0 | 6 | 0 | 0 | 6 | INF | 6 | | 6 | 3.13E-02 | 1.5 |
| Y0016 | chr18 | 5751556#5751576 | 5751576 | AT>CC_2 | Genotox-TS | | 5 0 | 5 | 0 | 0 | 5 | INF | 5 | 0 | 5 | 6.25E-02 | 1.2 |
| SY102 | chr18 | 5750933#5750967 | 5750933 | GA>AT_3 | Genotox-TS | | 1 4 | 1 | 4 | 1 | 4 | 4.0 | 5 | 1 | 5 | 3.75E-01 | 0.4 |

mutation # = SNP used when several available

**Notes**

P(two-tailed) = 2*BINOMDIST(n_low, n_low + n_high, 1/2, 1)

Monoclones (one allele = 0) with count ≥6 are statistically significant

statistically significant monoclone

HH, CPD hyperhotspot; HHW, highly mutated site adjacent to CPD hyperhotspot, in ETS-1 consensus binding site

Location colors indicate independent creation of the same mutation in more than one context or patient

P < 0.05

P < 0.001

-logP colors are a continuous gradient scale

### Table S3

### Table S4

**Table S4 Hybrid Capture Probe Sets**

**genome version: GRCh38/hg38**

**TwinStrand Probe Set 3**

| chr | Gene Name | Start | Stop | Probe Footprint (bp) | Reason To Select | Hyperhotspot nt |
|---|---|---|---|---|---|---|
| chr1 | *ARID1A* | 26763026 | 26763266 | 241 | Cancer driver reported mutated in sun-exposed skin (ref. 1) | - |
| chr1 | *ARID1A* | 26779001 | 26779327 | 327 | Cancer driver reported mutated in sun-exposed skin (ref. 1) | - |
| chr1 | *MROH7-TTC4//TTC4* | 54715766 | 54715946 | 181 | CPD Hyperhotspot | 54715855-56 |
| chr1 | *NOTCH2* | 119919494 | 119919674 | 181 | Cancer driver reported mutated in sun-exposed skin (ref. 1) | - |
| chr1 | *NOTCH2* | 119963670 | 119963850 | 181 | Cancer driver reported mutated in sun-exposed skin (ref. 1) | - |
| chr1 | *NOTCH2* | 119967424 | 119967604 | 181 | Cancer driver reported mutated in sun-exposed skin (ref. 1) | - |
| chr1 | *NOTCH2* | 119969541 | 119969721 | 181 | Cancer driver reported mutated in sun-exposed skin (ref. 1) | - |
| chr2 | *COQ10B* | 197453856 | 197454036 | 181 | CPD Hyperhotspot | 197453944-45 |
| chr3 | *DPH3//OXNAD1* | 16264909 | 16265089 | 181 | CPD Hyperhotspot | 16264996-97-98 |
| chr3 | *ULK4* | 41962151 | 41962331 | 181 | CPD Hyperhotspot | 41962240-41 |
| chr3 | *SMARCC1* | 47781835 | 47782015 | 181 | CPD Hyperhotspot | 47781923-24 |
| chr3 | *MST1R* | 49895717 | 49896317 | 601 | Candidate alternate splice site | - |
| chr3 | *RPL29* | 51995827 | 51996067 | 241 | CPD Hyperhotspot | 51995893-94, 51995943-44, 51995999-6000 |
| chr4 | *FGFR3* | 1802834 | 1803074 | 241 | Cancer driver reported mutated in sun-exposed skin (ref. 1) | - |
| chr4 | *FAT1* | 186599831 | 186600071 | 241 | Cancer driver reported mutated in sun-exposed skin (ref. 1) | - |
| chr5 | *ETF1* | 138543192 | 138543372 | 181 | CPD Hyperhotspot | 138543281-82 |
| chr5 | *RBM22* | 150701016 | 150701196 | 181 | CPD Hyperhotspot | 150701104-05 |
| chr7 | *EGFR* | 55143328 | 55143508 | 181 | Oncogene | - |
| chr7 | *EGFR* | 55174651 | 55174891 | 241 | Oncogene | - |
| chr7 | *BRAF* | 140753275 | 140753393 | 119 | Oncogene | - |
| chr8 | *COPS5//CSPP1* | 67062252 | 67062432 | 181 | CPD Hyperhotspot | 67062341-42 |
| chr9 | *PTCH1* | 95478981 | 95479221 | 241 | Mutated in BCC; Not detected in ref. 1 | - |
| chr9 | *PTCH1* | 95485649 | 95485889 | 241 | Mutated in BCC; Not detected in ref. 1 | - |
| chr9 | *NOTCH1* | 136499090 | 136499330 | 241 | Cancer driver reported mutated in sun-exposed skin (ref. 1) | - |
| chr9 | *NOTCH1* | 136500556 | 136500796 | 241 | Cancer driver reported mutated in sun-exposed skin (ref. 1) | - |
| chr9 | *NOTCH1* | 136502272 | 136502512 | 241 | Cancer driver reported mutated in sun-exposed skin (ref. 1) | - |
| chr9 | *NOTCH1* | 136517724 | 136517964 | 241 | Cancer driver reported mutated in sun-exposed skin (ref. 1) | - |
| chr11 | *ANAPC15* | 72112620 | 72112800 | 181 | CPD Hyperhotspot | 72112709-10 |
| chr11 | *ROI-3* | 77498452 | 77498632 | 181 | CPD Hyperhotspot | 77498541 |
| chr12 | *KRAS* | 25227162 | 25227462 | 301 | Oncogene | - |
| chr12 | *KRAS* | 25245214 | 25245514 | 301 | Oncogene | - |
| chr12 | *ROI-1* | 50840413 | 50840533 | 121 | CPD Hyperhotspot | 50840442 |
| chr15 | *TMEM87A//GANC* | 42273350 | 42273530 | 181 | CPD Hyperhotspot | 42273438-39 |
| chr15 | *CLPX* | 65185606 | 65185786 | 181 | CPD Hyperhotspot | 65185694-95 |
| chr17 | *TP53* | 7669499 | 7669799 | 301 | Tumor Suppressor | - |
| chr17 | *TP53* | 7670462 | 7670762 | 301 | Tumor Suppressor | - |
| chr17 | *TP53* | 7673139 | 7673929 | 791 | Tumor Suppressor | - |
| chr17 | *TP53* | 7674080 | 7674360 | 281 | Tumor Suppressor | - |
| chr17 | *TP53* | 7674755 | 7675324 | 570 | Tumor Suppressor | - |
| chr17 | *TP53* | 7675853 | 7676735 | 883 | Tumor Suppressor | - |
| chr17 | *TP53//LRG_321//RP11-199F11.2//WRAP53* | 7686070 | 7686250 | 181 | CPD Hyperhotspot | 7686158-59 |
| chr17 | *DCAF7* | 63550381 | 63550561 | 181 | CPD Hyperhotspot | 63550470-71 |
| chr18 | *Genotox-TS* | 5749264 | 5751664 | 2401 | TwinStrand Mutagenesis Region | - |
| chr19 | *NOTCH3* | 15161608 | 15161788 | 181 | Cancer driver reported mutated in sun-exposed skin (ref. 1) | - |
| chr19 | *NOTCH3* | 15192292 | 15192472 | 181 | Cancer driver reported mutated in sun-exposed skin (ref. 1) | - |
| chr22 | *ROI-6* | 46551374 | 46551554 | 181 | CPD Hyperhotspot | 46551463 |
| chrX | *RBM10* | 47185471 | 47185711 | 241 | Cancer driver reported mutated in sun-exposed skin (ref. 1) | - |
| chrX | *RBM10* | 47186101 | 47186341 | 241 | Cancer driver reported mutated in sun-exposed skin (ref. 1) | - |
| **Specific Genomic Regions Targeted:** | 48 | | | | | |
| **Total Probe Footprint:** | | | | 14413 | | |

**TwinStrand Probe Set 4**

| chr | Gene Name | Start | Stop | Probe Footprint (bp) | Reason To Select | Hyperhotspot nt |
|---|---|---|---|---|---|---|
| chr1 | *ARID1A* | 26763009 | 26763369 | 361 | Cancer driver reported mutated in sun-exposed skin (ref. 1) | - |
| chr1 | *ARID1A* | 26779001 | 26779394 | 394 | Cancer driver reported mutated in sun-exposed skin (ref. 1) | - |
| chr1 | *MROH7-TTC4//TTC4* | 54715766 | 54715946 | 181 | CPD Hyperhotspot | 54715855-56 |
| chr1 | *NOTCH2* | 119919462 | 119919702 | 241 | Cancer driver reported mutated in sun-exposed skin (ref. 1) | - |
| chr1 | *NOTCH2* | 119969447 | 119969807 | 361 | Cancer driver reported mutated in sun-exposed skin (ref. 1) | - |
| chr1 | *SCAMP3* | 155262203 | 155262383 | 181 | CPD Hyperhotspot | 155262293-94 |
| chr2 | *MRPL53//CCDC142* | 74472590 | 74472770 | 181 | CPD Hyperhotspot | 74472680-81 |
| chr2 | *COQ10B* | 197453856 | 197454036 | 181 | CPD Hyperhotspot | 197453944-45 |
| chr3 | *DPH3//OXNAD1* | 16264909 | 16265089 | 181 | CPD Hyperhotspot | 16264996-97-98 |
| chr3 | *ULK4* | 41962151 | 41962331 | 181 | CPD Hyperhotspot | 41962240-41 |
| chr3 | *MST1R* | 49895717 | 49896317 | 601 | Candidate alternate splice site | - |
| chr3 | *RPL29* | 51995827 | 51996067 | 241 | CPD Hyperhotspot | 51995893-94, 51995943-44, 51995999-6000 |
| chr3 | *UMPS* | 124730297 | 124730477 | 181 | CPD Hyperhotspot | 124730387-88 |
| chr3 | *RNA5SP146//RP11-550I24.2* | 157174902 | 157175082 | 181 | CPD Hyperhotspot | 157174992-93 |
| chr4 | *FGFR3* | 1802834 | 1803074 | 241 | Cancer driver reported mutated in sun-exposed skin (ref. 1) | - |
| chr4 | *FAT1* | 186599831 | 186600071 | 241 | Cancer driver reported mutated in sun-exposed skin (ref. 1) | - |
| chr5 | *ETF1* | 138543192 | 138543372 | 181 | CPD Hyperhotspot | 138543281-82 |
| chr5 | *RBM22* | 150701016 | 150701196 | 181 | CPD Hyperhotspot | 150701104-05 |
| chr6 | *TTK* | 80004429 | 80004609 | 181 | CPD Hyperhotspot | 80004518-19 |
| chr6 | *RMND1//C6orf211* | 151452156 | 151452336 | 181 | CPD Hyperhotspot | 151452245-46 |
| chr7 | *CYP3A5* | 99652751 | 99652931 | 181 | TwinStrand Microhaplotype Panel | - |
| chr8 | *COPS5//CSPP1* | 67062252 | 67062432 | 181 | CPD Hyperhotspot | 67062341-42 |
| chr9 | *NOTCH1* | 136499090 | 136499330 | 241 | Cancer driver reported mutated in sun-exposed skin (ref. 1) | - |
| chr9 | *NOTCH1* | 136500556 | 136500796 | 241 | Cancer driver reported mutated in sun-exposed skin (ref. 1) | - |
| chr9 | *NOTCH1* | 136502272 | 136502512 | 241 | Cancer driver reported mutated in sun-exposed skin (ref. 1) | - |
| chr10 | *CYP2C19* | 94761788 | 94761943 | 156 | TwinStrand Microhaplotype Panel | - |
| chr10 | *CYP2C8* | 95058235 | 95058475 | 241 | TwinStrand Microhaplotype Panel | - |
| chr10 | *CYP2C8* | 95067307 | 95067527 | 221 | TwinStrand Microhaplotype Panel | - |
| chr10 | *CYP2E1* | 133533319 | 133533544 | 226 | TwinStrand Microhaplotype Panel | - |
| chr10 | *CYP2E1* | 133537689 | 133537929 | 241 | TwinStrand Microhaplotype Panel | - |
| chr11 | *ANAPC15* | 72112620 | 72112800 | 181 | CPD Hyperhotspot | 72112709-10 |
| chr15 | *TMEM87A//GANC* | 42273350 | 42273530 | 181 | CPD Hyperhotspot | 42273438-39 |
| chr15 | *CLPX* | 65185606 | 65185786 | 181 | CPD Hyperhotspot | 65185694-95 |
| chr16 | *MC1R* | 89919607 | 89919847 | 241 | TwinStrand Microhaplotype Panel | - |
| chr17 | *TP53* | 7669499 | 7669799 | 301 | Tumor Suppressor | - |
| chr17 | *TP53* | 7670462 | 7670762 | 301 | Tumor Suppressor | - |
| chr17 | *TP53* | 7673139 | 7673929 | 791 | Tumor Suppressor | - |
| chr17 | *TP53* | 7674080 | 7674360 | 281 | Tumor Suppressor | - |
| chr17 | *TP53* | 7674755 | 7675324 | 570 | Tumor Suppressor | - |
| chr17 | *TP53* | 7675853 | 7676735 | 883 | Tumor Suppressor | - |
| chr17 | *TP53//LRG_321//RP11-199F11.2//WRAP53* | 7686070 | 7686250 | 181 | CPD Hyperhotspot | 7686158-59 |
| chr17 | *DCAF7* | 63550381 | 63550561 | 181 | CPD Hyperhotspot | 63550470-71 |
| chr18 | *Genotox-TS* | 5749264 | 5751664 | 2401 | TwinStrand Mutagenesis Region | - |
| chr19 | *DUS3L* | 5791092 | 5791272 | 181 | CPD Hyperhotspot | 5791181-82 |
| chrX | *RBM10* | 47185356 | 47185813 | 458 | Cancer driver reported mutated in sun-exposed skin (ref. 1) | - |
| chrX | *RBM10* | 47185998 | 47186382 | 385 | Cancer driver reported mutated in sun-exposed skin (ref. 1) | - |
| **Specific Genomic Regions Targeted:** | 47 | | | | | |
| **Total Probe Footprint:** | | | | 14721 | | |

**TwinStrand Probe Set 4 + *ADPRHL1***

| chr | Gene Name | Start | Stop | Probe Footprint (bp) | Reason To Select | Hyperhotspot nt |
|---|---|---|---|---|---|---|
| chr1 | *ARID1A* | 26763009 | 26763369 | 361 | Cancer driver reported mutated in sun-exposed skin (ref. 1) | - |
| chr1 | *ARID1A* | 26779001 | 26779394 | 394 | Cancer driver reported mutated in sun-exposed skin (ref. 1) | - |
| chr1 | *MROH7-TTC4//TTC4* | 54715766 | 54715946 | 181 | CPD Hyperhotspot | 54715855-56 |
| chr1 | *NOTCH2* | 119919462 | 119919702 | 241 | Cancer driver reported mutated in sun-exposed skin (ref. 1) | - |
| chr1 | *NOTCH2* | 119969447 | 119969807 | 361 | Cancer driver reported mutated in sun-exposed skin (ref. 1) | - |
| chr1 | *SCAMP3* | 155262203 | 155262383 | 181 | CPD Hyperhotspot | 155262293-94 |
| chr2 | *MRPL53//CCDC142* | 74472590 | 74472770 | 181 | CPD Hyperhotspot | 74472680-81 |
| chr2 | *COQ10B* | 197453856 | 197454036 | 181 | CPD Hyperhotspot | 197453944-45 |
| chr3 | *DPH3//OXNAD1* | 16264909 | 16265089 | 181 | CPD Hyperhotspot | 16264996-97-98 |
| chr3 | *ULK4* | 41962151 | 41962331 | 181 | CPD Hyperhotspot | 41962240-41 |
| chr3 | *MST1R* | 49895717 | 49896317 | 601 | Candidate alternate splice site | - |
| chr3 | *RPL29* | 51995827 | 51996067 | 241 | CPD Hyperhotspot | 51995893-94, 51995943-44, 51995999-6000 |
| chr3 | *UMPS* | 124730297 | 124730477 | 181 | CPD Hyperhotspot | 124730387-88 |
| chr3 | *RNA5SP146//RP11-550I24.2* | 157174902 | 157175082 | 181 | CPD Hyperhotspot | 157174992-93 |
| chr4 | *FGFR3* | 1802834 | 1803074 | 241 | Cancer driver reported mutated in sun-exposed skin (ref. 1) | - |
| chr4 | *FAT1* | 186599831 | 186600071 | 241 | Cancer driver reported mutated in sun-exposed skin (ref. 1) | - |
| chr5 | *ETF1* | 138543192 | 138543372 | 181 | CPD Hyperhotspot | 138543281-82 |
| chr5 | *RBM22* | 150701016 | 150701196 | 181 | CPD Hyperhotspot | 150701104-05 |
| chr6 | *TTK* | 80004429 | 80004609 | 181 | CPD Hyperhotspot | 80004518-19 |
| chr6 | *RMND1//C6orf211* | 151452156 | 151452336 | 181 | CPD Hyperhotspot | 151452245-46 |
| chr7 | *CYP3A5* | 99652751 | 99652931 | 181 | TwinStrand Microhaplotype Panel | - |
| chr8 | *COPS5//CSPP1* | 67062252 | 67062432 | 181 | CPD Hyperhotspot | 67062341-42 |
| chr9 | *NOTCH1* | 136499090 | 136499330 | 241 | Cancer driver reported mutated in sun-exposed skin (ref. 1) | - |
| chr9 | *NOTCH1* | 136500556 | 136500796 | 241 | Cancer driver reported mutated in sun-exposed skin (ref. 1) | - |
| chr9 | *NOTCH1* | 136502272 | 136502512 | 241 | Cancer driver reported mutated in sun-exposed skin (ref. 1) | - |
| chr10 | *CYP2C19* | 94761788 | 94761943 | 156 | TwinStrand Microhaplotype Panel | - |
| chr10 | *CYP2C8* | 95058235 | 95058475 | 241 | TwinStrand Microhaplotype Panel | - |
| chr10 | *CYP2C8* | 95067307 | 95067527 | 221 | TwinStrand Microhaplotype Panel | - |
| chr10 | *CYP2E1* | 133533319 | 133533544 | 226 | TwinStrand Microhaplotype Panel | - |
| chr10 | *CYP2E1* | 133537689 | 133537929 | 241 | TwinStrand Microhaplotype Panel | - |
| chr11 | *ANAPC15* | 72112620 | 72112800 | 181 | CPD Hyperhotspot | 72112709-10 |
| chr13 | *ADPRHL1* | 113449080 | 113449741 | 662 | CPD Hyperhotspot | 113449167-68, 113449201-02, 113449235-36, 113449302-03, 113449336-37, 113449370-71, 113449403-04, 113449471-72, 113449537-38, 113449571-72, 113449605-06 |
| chr15 | *TMEM87A//GANC* | 42273350 | 42273530 | 181 | CPD Hyperhotspot | 42273438-39 |
| chr15 | *CLPX* | 65185606 | 65185786 | 181 | CPD Hyperhotspot | 65185694-95 |
| chr16 | *MC1R* | 89919607 | 89919847 | 241 | TwinStrand Microhaplotype Panel | - |
| chr17 | *TP53* | 7669499 | 7669799 | 301 | Tumor Suppressor | - |
| chr17 | *TP53* | 7670462 | 7670762 | 301 | Tumor Suppressor | - |
| chr17 | *TP53* | 7673139 | 7673929 | 791 | Tumor Suppressor | - |
| chr17 | *TP53* | 7674080 | 7674360 | 281 | Tumor Suppressor | - |
| chr17 | *TP53* | 7674755 | 7675324 | 570 | Tumor Suppressor | - |
| chr17 | *TP53* | 7675853 | 7676735 | 883 | Tumor Suppressor | - |
| chr17 | *TP53//LRG_321//RP11-199F11.2//WRAP53* | 7686070 | 7686250 | 181 | CPD Hyperhotspot | 7686158-59 |
| chr17 | *DCAF7* | 63550381 | 63550561 | 181 | CPD Hyperhotspot | 63550470-71 |
| chr18 | *Genotox-TS* | 5749264 | 5751664 | 2401 | TwinStrand Mutagenesis Region | - |
| chr19 | *DUS3L* | 5791092 | 5791272 | 181 | CPD Hyperhotspot | 5791181-82 |
| chrX | *RBM10* | 47185356 | 47185813 | 458 | Cancer driver reported mutated in sun-exposed skin (ref. 1) | - |
| chrX | *RBM10* | 47185998 | 47186382 | 385 | Cancer driver reported mutated in sun-exposed skin (ref. 1) | - |
| **Total Probe Footprint:** | | | | 15383 | | |

**TwinStrand Probe Set 5**

| chr | Gene Name | Start | Stop | Probe Footprint (bp) | Reason To Select | Hyperhotspot nt |
|---|---|---|---|---|---|---|
| chr1 | *ARID1A* | 26763009 | 26763369 | 361 | Cancer driver reported mutated in sun-exposed skin (ref. 1) | - |
| chr1 | *ARID1A* | 26779001 | 26779394 | 394 | Cancer driver reported mutated in sun-exposed skin (ref. 1) | - |
| chr1 | *MROH7-TTC4//TTC4* | 54715766 | 54715946 | 181 | CPD Hyperhotspot | 54715855-56 |
| chr1 | *NOTCH2* | 119919462 | 119919702 | 241 | Cancer driver reported mutated in sun-exposed skin (ref. 1) | - |
| chr1 | *NOTCH2* | 119969447 | 119969807 | 361 | Cancer driver reported mutated in sun-exposed skin (ref. 1) | - |
| chr1 | *SCAMP3* | 155262203 | 155262383 | 181 | CPD Hyperhotspot | 155262293-94 |
| chr2 | *MRPL53//CCDC142* | 74472590 | 74472770 | 181 | CPD Hyperhotspot | 74472680-81 |
| chr2 | *COQ10B* | 197453856 | 197454036 | 181 | CPD Hyperhotspot | 197453944-45 |
| chr3 | *DPH3//OXNAD1* | 16264909 | 16265089 | 181 | CPD Hyperhotspot | 16264996-97-98 |
| chr3 | *ULK4* | 41962151 | 41962331 | 181 | CPD Hyperhotspot | 41962240-41 |
| chr3 | *RPL29* | 51995827 | 51996067 | 241 | CPD Hyperhotspot | 51995893-94, 51995943-44, 51995999-6000 |
| chr3 | *UMPS* | 124730297 | 124730477 | 181 | CPD Hyperhotspot | 124730387-88 |
| chr3 | *RNA5SP146//RP11-550I24.2* | 157174902 | 157175082 | 181 | CPD Hyperhotspot | 157174992-93 |
| chr5 | *CENPH* | 69189438 | 69189618 | 181 | CPD Hyperhotspot | 69189528-29 |
| chr5 | *ETF1* | 138543192 | 138543372 | 181 | CPD Hyperhotspot | 138543281-82 |
| chr5 | *RBM22* | 150701016 | 150701196 | 181 | CPD Hyperhotspot | 150701104-05 |
| chr6 | *TTK* | 80004429 | 80004609 | 181 | CPD Hyperhotspot | 80004518-19 |
| chr6 | *PPIL4* | 149546000 | 149546300 | 301 | CPD Hyperhotspot | 149546149-50 |
| chr6 | *RMND1//C6orf211* | 151452156 | 151452336 | 181 | CPD Hyperhotspot | 151452245-46 |
| chr8 | *COPS5//CSPP1* | 67062252 | 67062432 | 181 | CPD Hyperhotspot | 67062341-42 |
| chr9 | *URM1* | 128371263 | 128371443 | 181 | CPD Hyperhotspot | 128371292-93 |
| chr9 | *NOTCH1* | 136499090 | 136499330 | 241 | Cancer driver reported mutated in sun-exposed skin (ref. 1) | - |
| chr9 | *NOTCH1* | 136500556 | 136500796 | 241 | Cancer driver reported mutated in sun-exposed skin (ref. 1) | - |
| chr9 | *NOTCH1* | 136502272 | 136502512 | 241 | Cancer driver reported mutated in sun-exposed skin (ref. 1) | - |
| chr10 | *RP11-80H5.9//KIF20B* | 89701414 | 89701594 | 181 | CPD Hyperhotspot | 89701503-04 |
| chr10 | *CYP2C19* | 94761788 | 94761943 | 156 | TwinStrand Microhaplotype Panel | - |
| chr11 | *ANAPC15* | 72112620 | 72112800 | 181 | CPD Hyperhotspot | 72112709-10 |
| chr12 | *WBP11//C12orf60* | 14803498 | 14803678 | 181 | CPD Hyperhotspot | 14803587-88 |
| chr13 | *MRPS31* | 40771120 | 40771240 | 121 | CPD Hyperhotspot | 40771209-10 |
| chr13 | *ADPRHL1* | 113449080 | 113449741 | 662 | CPD Hyperhotspot | 113449167-68, 113449201-02, 113449235-36, 113449302-03, 113449336-37, 113449370-71, 113449403-04, 113449471-72, 113449537-38, 113449571-72, 113449605-06 |
| chr15 | *TMEM87A//GANC* | 42273350 | 42273530 | 181 | CPD Hyperhotspot | 42273438-39 |
| chr15 | *CLPX* | 65185606 | 65185786 | 181 | CPD Hyperhotspot | 65185694-95 |
| chr17 | *TP53* | 7669499 | 7669799 | 301 | Tumor Suppressor | - |
| chr17 | *TP53* | 7670462 | 7670762 | 301 | Tumor Suppressor | - |
| chr17 | *TP53* | 7673139 | 7674360 | 1222 | Tumor Suppressor | - |
| chr17 | *TP53* | 7674755 | 7675324 | 570 | Tumor Suppressor | - |
| chr17 | *TP53* | 7675853 | 7676492 | 640 | Tumor Suppressor | - |
| chr17 | *TP53* | 7676495 | 7676735 | 241 | Tumor Suppressor | - |
| chr17 | *TP53//LRG_321//RP11-199F11.2//WRAP53* | 7686070 | 7686250 | 181 | CPD Hyperhotspot | 7686158-59 |
| chr17 | *DCAF7* | 63550381 | 63550561 | 181 | CPD Hyperhotspot | 63550470-71 |
| chr18 | *Genotox-TS* | 5749264 | 5751664 | 2401 | TwinStrand Mutagenesis Region | - |
| chr19 | *DUS3L* | 5791092 | 5791272 | 181 | CPD Hyperhotspot | 5791181-82 |
| chr22 | *NOL12* | 37686415 | 37686655 | 241 | CPD Hyperhotspot | 37686564-65 |
| **Specific Genomic Regions Targeted:** | 43 | | | | | |
| **Total Probe Footprint:** | | | | 13641 | | |

**NOTES:**

Samples using TwinStrand Probe Set 3:
**FS2A, FS2B, FS3A, FS3B, Saliva 1, Saliva 2, Saliva 3, Saliva 4, YUMIPS, YUTENT, YUDOBI**

Samples using TwinStrand Probe Set 4:
**DTC, YUMIPS, YUTENT, YUDOBI, Chronic UV Experiment**

Samples using TwinStrand Probe Set 4 + *ADPRHL1*:
**FS3A rerun, FS3B rerun, DTC, DNA-63, Y008CTCL, Y0011CTCL, SY102CTCL, SY107CTCL, YUTENT**

Samples using TwinStrand Probe Set 5:
**DTC, SY114CTCL, SY118CTCL, SY112TPN, Y0015AM, Y0016AM, Y0017AM, SY132AM, SY146AM, SY124AM, SY126AM, SY130AM, SY138AM, UM001AM, UM002AM, UM003AM, UM005AM**

| Term | Definition |
| --- | --- |
| Duplex Consensus Sequence | One DNA sequence embodying consensus between the top and bottom strands of a single duplex molecule. Obtained from PCR copies of differentially tagged top and bottom strands of the original duplex molecule. |
| DCS | A duplex consensus sequence derived by a specific bioinformatics procedure that includes realignment of duplex consensus sequences after construction from single-strand consensus sequences.<br>References: [30, 64] |
| IC-DCS | IndelCorrected Duplex Consensus Sequence. A duplex consensus sequence derived from separately sequenced and correctly one-to-one genome-aligned PCR copies of the differentially tagged top and bottom strands of the original duplex molecule. |
| Strand-specific UMI | The sequencing library is constructed to attach a different unique molecular identifier sequence (UMI) to the top and bottom strands of each duplex molecule pre-PCR. |
| Correctly structured read pair | Has a different error-free unique molecular identifier sequence (UMI) at each end, termed the ditag. |
| Contiguous region of interest (CRI) | Double-stranded genome region spanned by a single-strand capture probe. For alignment purposes, it also includes 250 bp of flanking sequence. |

| | |
|---|---|
| onTarget | Mapped into the particular probe's CRI plus 250 bp flanks. |
| onTarget ditag read family | All correctly structured onTarget read pairs that share the same ditag, start nt, end nt, and CIGAR string. Both strands must be included for an acceptable family. This is merely a precise statement of "a read family". |
| Single-end IC single-strand consensus sequence | For one end of the library insert, a consensus sequence derived from $\geq$1 PCR progeny of one parental strand in an accepted onTarget ditag read family. |
| Single-end IC-DCS | For one end of the library insert, a consensus sequence derived from a pair of single-end IC single-strand consensus sequences. |
| Paired-end IC-DCS | A pair of single-end IC-DCSs. It is thus constructed from a family of >2 PCR progeny and having both strands of a single parental paired-end duplex, with each family member consisting of a correctly structured read pair sharing the same start, end, and CIGAR string and having unique full-length mapping onto the probe's contiguous region of interest (including 250 bp flanks). Every overlapping pair of single-end IC-DCSs is replaced by a single consensus IC-DCS. |
| **Two Estimates of Family Sizes** | |
| Average family size = (correctly structured read pairs uniquely mapped onto a CRI in chrN) / (onTarget ditag read families) This parameter aids in deciding whether the sequencing run size suffices to get the desired coverage of a CRI. | |
| Average useful-family size = (correctly structured read pairs uniquely mapped onto a CRI in chrN) / (onTarget ditag read families that have size >2 and have both strands) This parameter aids in describing the useful families; the non-useful families that aren't included reduce the coverage for that CRI. | |

**Table S5. Definition of terms in IndelCorrected-DCS (IC-DCS) alignment pipeline.**

| | 0 J/m² | | | 300 J/m² | | | 1000 J/m² | | |
|---|---|---|---|---|---|---|---|---|---|
| Irradiation | Days after 1st irradiation | Collection timepoint (days after prior irradiation) | Collection timepoint (days after first irradiation) | Days after 1st irradiation | Collection timepoint (days after prior irradiation) | Collection timepoint (days after first irradiation) | Days after 1st irradiation | Collection timepoint (days after prior irradiation) | Collection timepoint (days after first irradiation) |
| 1st | 0 | 1 | 1 | 0 | 1 | 1 | 0 | 1 | 1 |
| 2nd | 3 | 1 | 4 | 3 | 1 | 4 | 3 | 1 | 4 |
| 3rd | 6 | 2 | 8 | 6 | 2 | 8 | 6 | 2 | 8 |
| 4th | 10 | 2 | 12 | 10 | 2 | 12 | 10 | 3 | 13 |
| 5th | 15 | 2 | 17 | 15 | 2 | 17 | 15 | 4 | 19 |
| 6th | 19 | 2 | 21 | 19 | 2 | 21 | 26 | 4 | 30 |
| 7th | 24 | 2 | 26 | 24 | 4 | 28 | | | |
| 8th | 30 | 3 | 33 | 30 | 3 | 33 | | | |

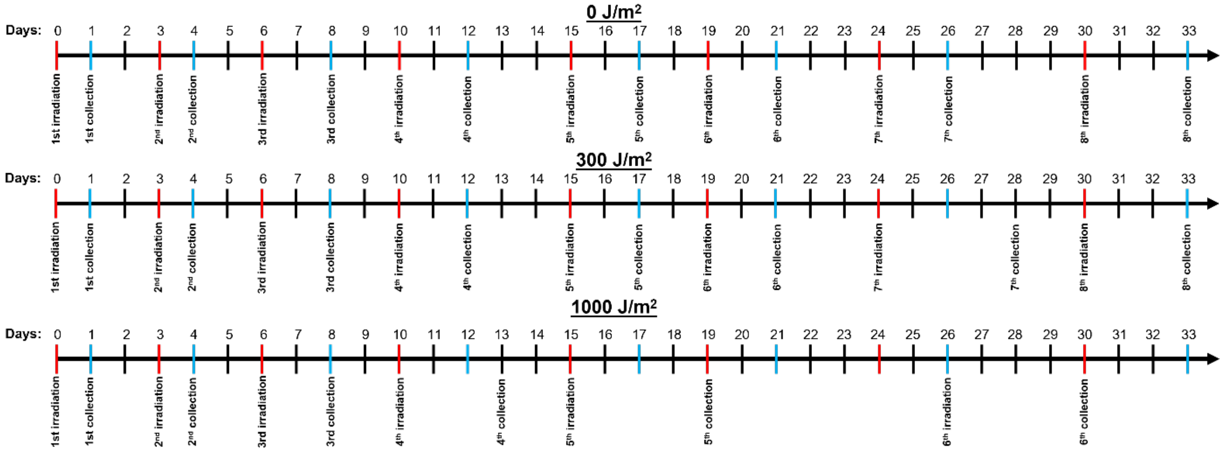

**Fig. S1. Chronic mutation timeline.**

#### Supplementary Results

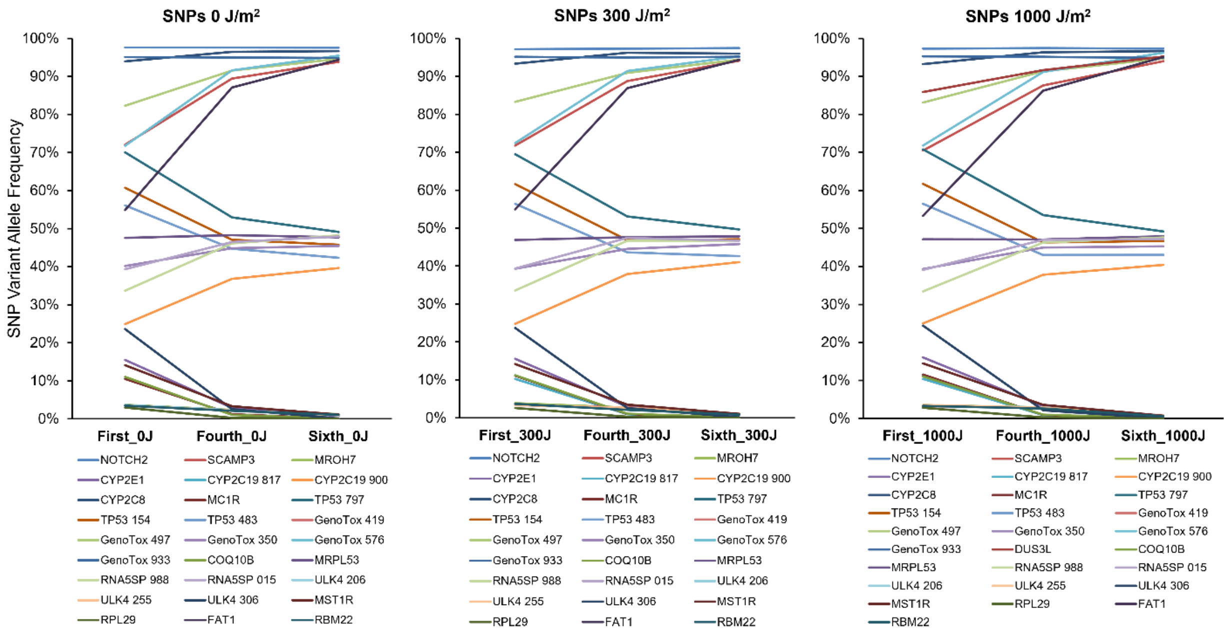

**Fig. S2. Selection against particular donors revealed by *in vitro* reductions in VAF of single-nucleotide polymorphisms (SNPs).**
